# Recognition and catalytic mechanism of tRNA m^3^C methyltransferase METTL6

**DOI:** 10.1101/2025.06.25.660299

**Authors:** Jana Aupič, Iven Hollmer, Laura Tengo, Peter Sehr, Eva Kowalinski, Alessandra Magistrato

## Abstract

Post-transcriptional RNA modifications regulate the RNA structural and functional landscape, and thus play a central role in many cellular processes. Accordingly, aberrant RNA modification pathways contribute to a variety of human diseases, including cancer and neurological disorders. However, recognition and catalytic mechanisms of most RNA-modifying proteins remain obscure, due to sparse structural data. Recently, the cryogenic electron microscopy structure of N3-methyl-cytosine (m^3^C) methyltransferase METTL6 revealed seryl-tRNA synthetase (SerRS) as a cofactor for tRNA^Ser^ selection, but left the cooperative role of SerRS and the reaction process undefined. Here, combining computer simulations and *in vitro* experiments, we define the mechanism of concerted substrate identification by METTL6 and SerRS, demonstrating that SerRS pre-organizes the tRNA^Ser^ variable arm for METTL6 recruitment. We show that once METTL6 is stably docked onto the SerRS-tRNA^Ser^ complex, thereby ensuring the precise alignment of the reactants, the methylation reaction proceeds spontaneously. Our results establish the catalytic mechanism of m^3^C methyltransferases and provide the foundation for rational design of targeted therapeutics.

## Introduction

RNA molecules possess remarkable structural and functional diversity, rivalling that of proteins. While composed of only four nucleotides, the chemical richness of RNA is greatly enhanced by post-transcriptional modifications, with more than 170 modifications uncovered so far ^1–4^. Among the different RNA types, tRNA molecules have the highest density of RNA modifications ^5^. RNA modifications influence stability, folding, structure and function of RNA molecules and therefore contribute to the regulation of gene expression ^1,6–10^. For tRNA molecules this is particularly evident, since covalent chemical modifications affect their function as mRNA decoders in protein synthesis ^11–13^. Modifications in the anticodon stem loop directly affect tRNA-mRNA codon pairing at the ribosome and thus influence translation fidelity (i.e. correct reading of the codon sequence) and translation rate ^13–20^. On the other hand, modifications in regions outside of the anticodon stem loop regulate mRNA translation by modulating tRNA localization, tRNA stability and tRNA folding, which can affect its interactions with the appropriate aminoacyl-tRNA synthetase and/or the ribosome ^13,21–25^. Thus, deposition or removal of tRNA modifications can lead to preferential translation of mRNA transcripts with high content of certain codons ^16,25–28^. Several lines of evidence suggest that this so-called codon-biased translation is an important mechanism of cellular response to changing environmental factors, such as high oxidative stress ^19,28–31^. Additionally, cancer cells were also shown to employ codon-bias for proteome reprogramming to drive cancer cell phenotypes, such as cell proliferation, metastasis and cell survival ^13,32–36^.

Methylation is the most frequent RNA modification. It is catalyzed by a diverse group of RNA methyltransferases (RNA-MTases). These enzymes transfer a methyl group from a small molecule cofactor, most frequently S-Adenosyl-methionine (SAM), to their substrate ^37,38^. Most RNA-MTases carry a conserved Rossmann-fold core coordinating the methyl-donor SAM ^37,39^. In addition, they typically include auxiliary domains for RNA binding or interact with external factors to enhance their affinity and specificity for a particular substrate ^37,39^. Transient interactions between RNA-MTases and their substrates complicate high-resolution structural studies, limiting insights into substrate recognition and catalysis of many RNA-MTases.

The N3-methylcytidine (m^3^C) modification has recently received increased attention due to its clear role in ensuring translation fidelity ^21,40^. m^3^C is present at position 32 in the anticodon loop of selected tRNAs in eukaryotes ^21,40–42^. While the yeast enzymes catalyzing m^3^C_32_ formation seem multifunctional ^43–46^, in humans three enzymes, METTL6, along with METTL2 and METTL8, are the tRNA-MTases responsible for the deposition of m^3^C_32_ in specific tRNA substrates ^40,47–51^. In particular, METTL6 catalyzes the formation of m^3^C in the anticodon stem loop of all serine isoacceptors (tRNA^Ser^) ^40,48^, while METTL2 works on cytosolic tRNA^Thr^ and tRNA^Arg^ ^47,48^. METTL8, located in the mitochondria, mediates the methylation of mitochondrial (mt) tRNA^Thr^ and tRNA^Ser^ ^47,49–51^. All m^3^C-tRNA-MTases utilize SAM as the source of the methyl group and display remarkable sequence and structure similarity ^41,52,53^. Several studies have shown that the activity of METTL6 and related yeast enzymes targeting tRNA^Ser^ is strongly dependent on the presence of seryl-tRNA synthetase (SerRS) ^43,47,54^. SerRS is a vital and highly abundant enzyme responsible for the attachment of the serine aminoacyl moiety to all seryl-tRNAs ^55,56^. Recently, we have revealed through biochemical assays and cryogenic electron microscopy (cryo-EM) how the tRNA-MTase METTL6 and SerRS cooperate to achieve efficient and specific tRNA^Ser^ recognition ^54^. The METTL6-SerRS-tRNA^Ser^ cryo-EM structure demonstrated that SerRS binds the tRNA^Ser^ acceptor arm with the aminoacylation site of the tRNA^Ser^ in its active site. Previous works showed that since serine tRNAs are diverse in their anticodon sequences (AGA, CGA, UGA, GCU), SerRS utilizes the long variable arm of serine tRNAs as the identity element for specific recognition ^57–59^. Indeed, in the METTL6-SerRS-tRNA^Ser^ complex structure, the long variable arm is sandwiched between SerRS and METTL6. Although our cryo-EM structure identified the interactions formed between METTL6 and SerRS, the mechanism by which SerRS increases the reaction efficiency of METTL6 remained unclear. Furthermore, while our structural data revealed the tripartite tRNA recognition domain present in all m^3^C-tRNA-MTases ^54^, the exact progression of the catalytic mechanism of this enzyme family remained unknown, due to the lack of identifiable catalytic residues.

Attaining a comprehensive understanding of the m^3^C-tRNA-MTases functional mechanism is particularly pertinent for the development of novel therapeutic agents as all m^3^C-tRNA-MTases are indicated to be relevant in different diseases. In particular, METTL6 expression is deregulated in multiple cancer types, including pancreatic, breast and liver cancer ^40,60–62^. Knockout of METTL6 in mouse embryonic stem cells reduced liver cancer cell proliferation and impaired colony formation capacity, suggesting it is a viable anticancer target ^40^. Furthermore, breast and lung cancer cells lacking METTL6 showed greater sensitivity to cisplatin ^21,63^.

To resolve the mechanism of how SerRS facilitates tRNA^Ser^ recognition by METTL6 and to reveal the details of m^3^C catalysis, we herein combine classical and quantum-classical molecular dynamics (MD) simulations. We challenge our computational results using in vitro biochemical and biophysical assays of different METTL6 mutants. Through this integrative approach, we uncover that SerRS enhances allosteric communication between the anticodon stem loop binding site and the N-terminal domain of METTL6. This interdomain coupling facilitates the accurate docking of the tRNA methyl acceptor residue C^32^ into the METTL6 active site, positioning it adjacent to the methyl donor SAM. This induced proximity of methyl-donor and acceptor then propels the spontaneous catalysis, leading to highly efficient and specific methylation of C_32_. The disclosed mechanism likely applies to all other m^3^C-tRNA-MTases.

## Results

### Reconstruction of the pre-catalytic state of the METTL6-SerRS-tRNA^Ser^ complex

Previous data indicated that adding SerRS to a methylation assay significantly increased METTL6 activity ^43,47,54^. To resolve the underlying mechanism for this enhancement, we performed classical MD simulations to assess the role of SerRS for tRNA^Ser^ recognition by METTL6. SerRS forms an obligatory dimer, able to interact with one or two tRNAs ^47,56^. While theoretically two METTL6 molecules could independently access these tRNAs, the lower abundance of METTL6 in the cell compared to SerRS ^64^, and the fact that the tRNA molecules in the cryo-EM structure of the METTL6–SerRS–tRNA^Ser^ complex do not communicate, suggest that the functional m^3^C_32_ methylation unit is composed of the SerRS dimer, one molecule of tRNA^Ser^ and one copy of METTL6. Hence, we initiated our computer simulations from the METTL6-SerRS-tRNA^Ser^ complex that contains a single METTL6 as this was resolved at the highest resolution (PDBID: 8P7B, Figure 1a) ^54^.

**Figure 1.**
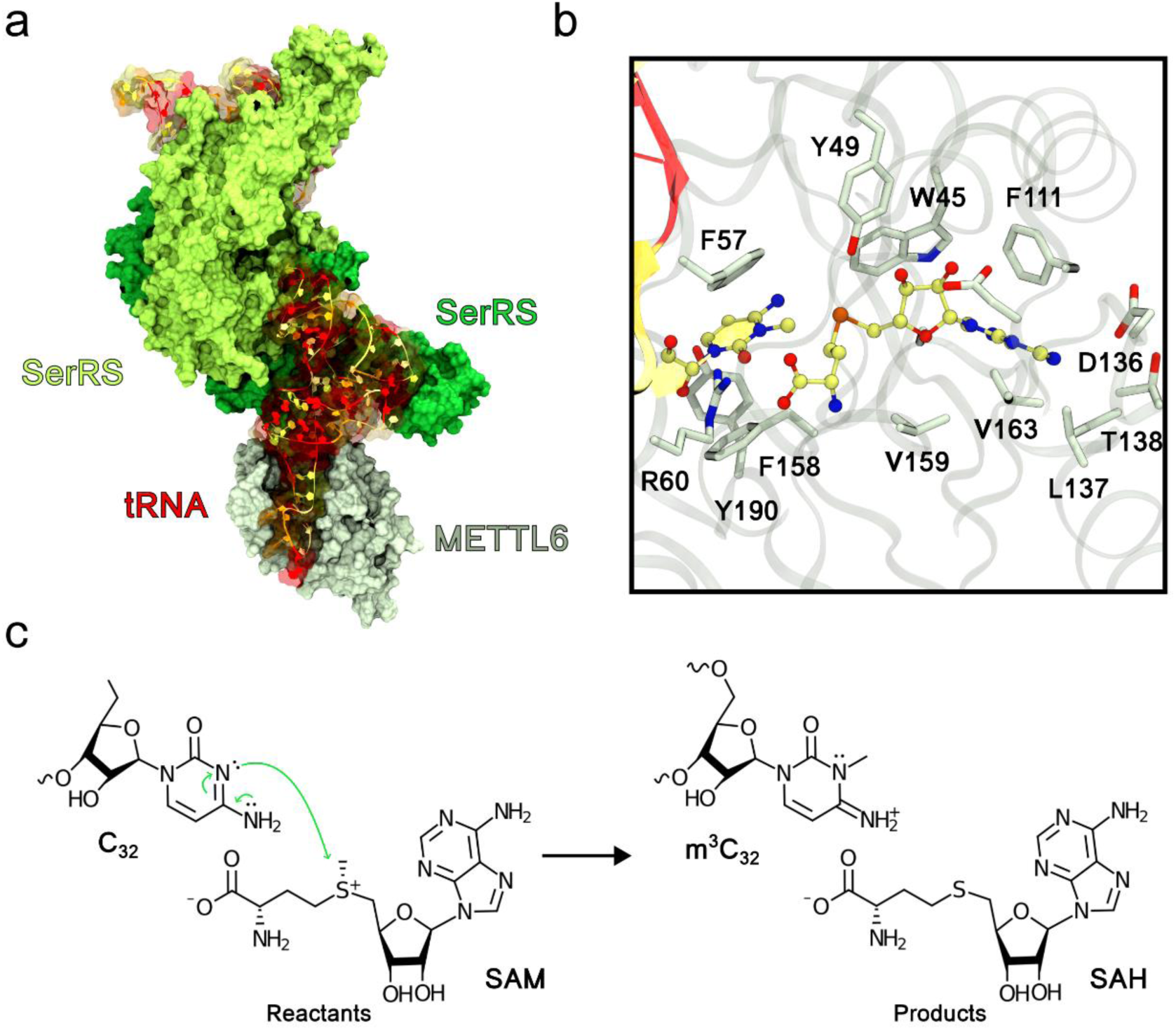
METTL6 is a SAM-based m^3^C methyltransferase. (a) Cryo-EM structure of the 1:2:2 METTL6-SerRS-tRNA^Ser^ complex. SerRS and METTL6 are shown in green and white surface representation, respectively, and tRNA^Ser^ is shown in ribbon representation. (b) Close up view of the catalytic site of METTL6 in the product state as observed in the cryo-EM structure. Residues participating in m^3^C_32_ and SAH binding are shown as licorice. (c) Scheme of the methylation reaction. The free electron pair on the N3 atom of C_32_ carries out a nucleophilic attack on the methyl group of SAM, bound to the positively charged sulfonium cation. This results in the formation of the positively charged m^3^C_32_ and neutral SAH.

In the cryo-EM sample, the active site of METTL6 was trapped in the post-methylation product state, with the active site containing the m^3^C_32_ and demethylated SAM, i.e. the S-adenosyl-L-homocysteine (SAH) product (Figures 1b, c). In the cryo-EM structure, SAH forms hydrogen bonds to Tyr49, Arg60 and Asp110 with its amino acid part and ribose ring. Additionally, the SAH adenosyl moiety is anchored by forming hydrophobic interactions with Phe111, Leu137, Tyr138, Ala162 and Val163 (Figure 1b). On the other hand, the methyl acceptor base, m^3^C_32_, was observed in the active site in a flipped-out conformation, oriented away from the interior of the anticodon stem loop. The m^3^C_32_ orientation is stabilized by base stacking to Phe57 and Phe158. Additionally, Arg60 forms a hydrogen bond with the C_32_ nucleobase carbonyl group.

For our computer simulations, we modified the active site coordinates from the product state to the reactant (pre-methylation) state, replacing m³C_32_ with C_32_ and SAH with SAM. To achieve this, we broke the covalent bond connecting the C_32_:N3 atom and the methyl group, while reconnecting the latter to the SAH:S atom. To dissect the long-range implications of SerRS, we conducted classical MD simulations of two systems: the full METTL6-SerRS-tRNA^Ser^ complex, consisting of two SerRS copies, two tRNA^Ser^ molecules, and a single METTL6, and the METTL6-tRNA^Ser^ complex without SerRS. Both systems were described with the CHARMM36m force field ^65^. The complexes were solvated with TIP3P water and surrounded with KCl ions at 150 mM concentration. Simulations were run in Gromacs for 500 ns in triplicates ^66^.

### Seryl-tRNA synthetase impacts tRNA recognition through interaction with the METTL6 N-terminal helices and insertion motif

Overall, the METTL6-SerRS-tRNA^Ser^ complex remained stably folded during classical MD simulations, with all components largely preserving their native interfacial contacts (Figures 2a-e, Fig. S1). Additionally, C_32_ and SAM remained stably bound in the active site (Figs. S2 and S3), with their relative positioning conducive for the methyltransferase reaction in 72 ± 18 % of simulation frames (Figure 2d, Fig. S4). To determine which molecular features are most important for allosteric communication between tRNA^Ser^ and METTL6 we performed dynamical network analysis using generalized correlations (Figures 2a-c) ^67^. Briefly, based on inter-residue contacts, dynamical network analysis allows us to determine regions of proteins, so-called communities, that move in concert and show a high degree of correlation. Often these communities correspond or overlap with a topological region of the protein or the nucleic acid molecule. Each pair of residues is connected by a so-called edge. The importance of each edge for communication corresponds to its betweenness value, which equals the number of shortest paths between all residue pairs that pass through that edge. Thus, edges characterized by high betweenness values are the most important for communication within the complex. By determining how information is exchanged along the edges, it is possible to identify the dominant communication channels within molecules.

**Figure 2.**
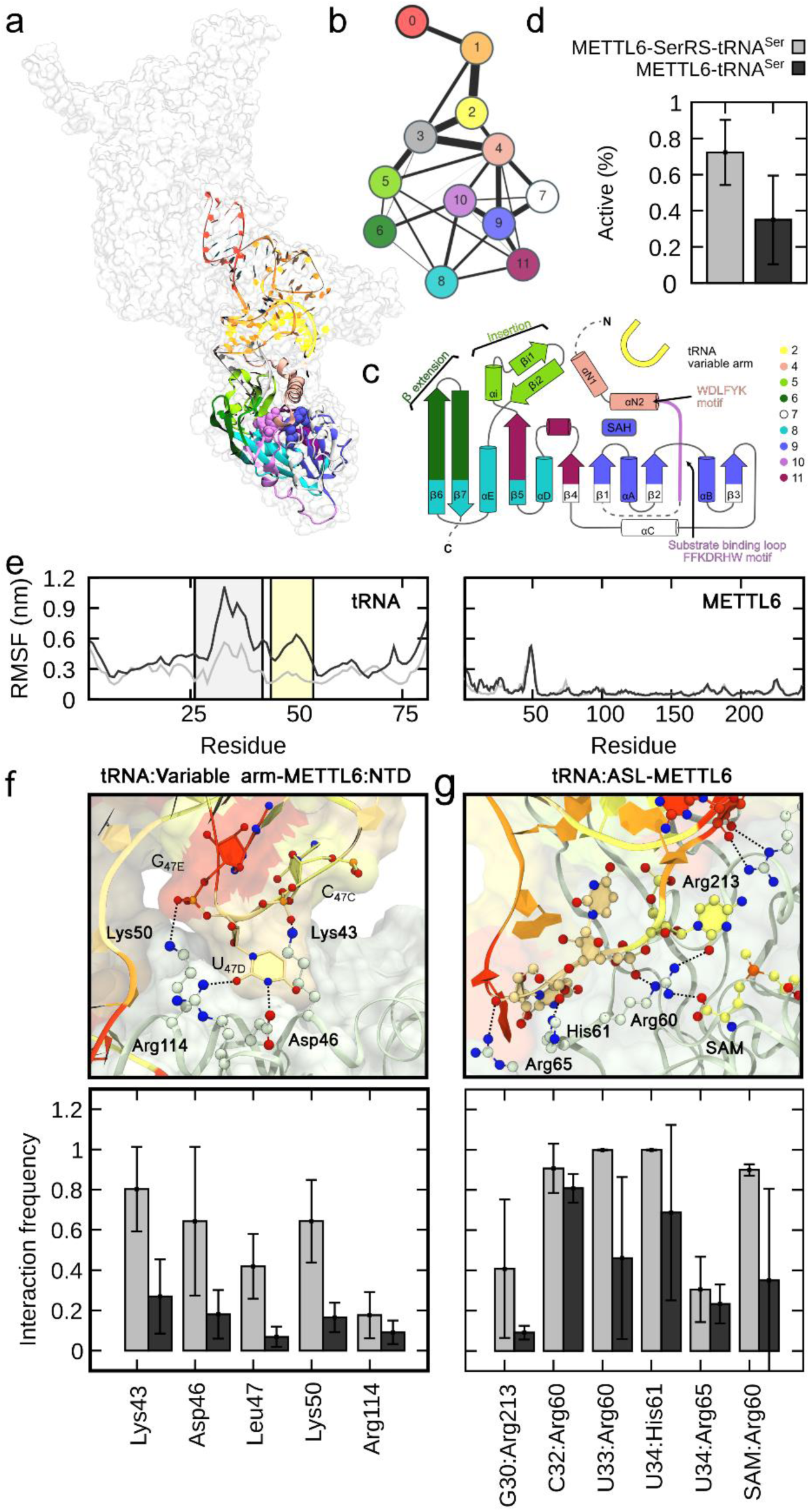
SerRS interactions with tRNA^Ser^ long variable arm and METTL6 N-terminal domain facilitate tRNA^Ser^ recognition by METTL6. (a) Dynamical network analysis of communication pathways between METTL6 and tRNA^Ser^ in the full 1:2:2 METTL6-SerRS-tRNA^Ser^ complex. Colors indicate different communities. (b) A simplified, coarse schematic representation of the METTL6-SerRS-tRNA^Ser^ communication network. Circles represent individual communities and edges are depicted with black lines. Edge thickness corresponds to its betweenness value. (c) The Rossmann fold scheme of METTL6 colored according to community analysis. (d) Probability of a reactive conformation of the active site as observed from classical molecular dynamics. (e) Average residue root mean square fluctuations (RMSF) for tRNA^Ser^ (left panel) and METTL6 (right panel). In the left panel, the areas shaded in gray and yellow correspond to the anticodon stem loop and the variable arm, respectively. The light gray line reports results for the full complex, while the dark gray line depicts results obtained for the METTL6-tRNA^Ser^ complex. (f) Depiction of hydrogen bonding between METTL6 N-terminal domain (NTD) and tRNA^Ser^ variable arm. The bottom panel reports interaction frequencies. Light gray bars correspond to the full complex, while dark gray bars show results for the METTL6-tRNA^Ser^ complex. (g) Depiction of hydrogen bonding between METTL6 and tRNA^Ser^ anticodon stem loop (ASL). Bottom panel reports interaction frequencies. Gray bars correspond to the full complex, while dark gray bars show results for the METTL6-tRNA^Ser^ complex. Error bars denote ± s.d. from the mean for *n* = 3 simulation replicas.

The tRNA binding domain of m^3^C-tRNA-MTases (m^3^C-RBD) is composed of three short motifs interlaced within the central Rossmann-fold: two N-terminal helices that are followed by the substrate binding loop containing the FFKDRHW motif conserved across all m^3^C-tRNA-MTases; an insertion motif between strand β5 and helix αE; and a β-extension of the C-terminal strands β6 and β7 ^54^. The analysis of our MD trajectories collected for the METTL6-SerRS-tRNA^Ser^ complex revealed that the binding of the tRNA^Ser^ anticodon stem (community 3 in Figures 2a-c) primarily occurs via the m^3^C-RBD insertion motif (residues 195-220, community 5). Conversely, the tRNA^Ser^ anticodon loop (communities 6 and 10) displayed tight interaction with the substrate binding loop (residues 55-62, community 10) and the β-extension region (residues 246-259, community 6). Further, our simulations indicated that the two N-terminal helices (ɑN1 and ɑN2, community 4) of METTL6 contribute significantly to anticodon stem loop docking at the active site. The influence of these ɑ helices was propagated both directly (edge between communities 3 and 4 or 10 and 4) and indirectly through the tRNA^Ser^ variable arm (community 2), driven by interactions with charged METTL6 residues (Lys43, Asp46, Lys50 and Arg51) all located on helix ɑN2. These residues formed hydrogen bonds to the RNA backbone and the nucleobase of U^47D^, which is located at the extremity of the serine-specific long variable arm. Taken together, our data identify the N-terminal helices, the insertion motif and the β-extension region of METTL6 as key structural elements for docking of the tRNA^Ser^ anticodon stem loop and, additionally, highlight the importance of tight interactions between the N-terminal helices and the tRNA^Ser^ variable arm. This is particularly notable, as SerRS primarily interacts directly with the METTL6 N-terminal helices and, secondly, the long variable arm is a distinctive feature of serine tRNA isoacceptors, which SerRS uses for substrate selection ^68,69^.

### Seryl-tRNA synthetase facilitates tRNA^Ser^ long variable arm binding to METTL6

Strikingly, in MD simulations of the minimal METTL6-tRNA^Ser^ complex (Fig. S5 to S8) – performed in the absence of SerRS - the interactions between the tRNA^Ser^ variable arm and helix ɑN2 were significantly disrupted, leading to a complete detachment of the tRNA^Ser^ variable arm from the helix ɑN2 (Figure 2f). This resulted in an overall increased flexibility of the variable arm with long-range effects provoking the destabilization of the anticodon stem loop secondary structure (Figure 2e, Fig. S6). In the absence of SerRS, the interaction between the anticodon stem loop and METTL6 was markedly destabilized, as evidenced by the melting of hydrogen bonds between the RNA phosphodiester backbone near the methylation target C_32_ and positively charged residues of METTL6 (Arg60, His61 and Arg 213) (Figure 2g, Fig. S7). The loss of these interactions led to the displacement of C_32_ from the catalytic site (Fig. S8), starkly reducing the frequency of catalytically competent active site conformations (from 72 ± 18 % to 35 ± 24 %, Figure 2d). In conclusion, our MD simulations reveal mechanistic insights into the role of SerRS as a cofactor of METTL6 in recognizing and binding tRNA^Ser^. SerRS facilitates the correct positioning of the anticodon stem loop, including the flipped-out base C_32_, into the METTL6 active site, making it accessible for modification.

To evaluate whether the outcomes of our MD simulations were affected by force field selection, we repeated the simulations of the minimal METTL6-tRNA^Ser^ complex with a different force field (i.e. Amber force field, see Methods). These new simulations confirmed an increased flexibility of the tRNA^Ser^ variable arm and anticodon loop and the weakening of interactions between tRNA^Ser^ and METTL6 (Figs. S9 to S11). However, detachment of C_32_ from the catalytic cavity was rarer and observed only transiently (Fig. S12). Thus, both Amber and Charmm force fields indicate that the contribution of SerRS is crucial for the stability of the METTL6-tRNA^Ser^ complex and vital for the appropriate positioning of the reactive groups in the catalytic cavity.

### The N-terminal helix *α*N2 is an allosteric signal transducer within the METTL6-SerRS-tRNA^Ser^ complex

Our computer simulations indicate that SerRS facilitates the binding of the tRNA^Ser^ long variable arm to METTL6. Therefore, we probed the interface between the variable arm of tRNA^Ser^ and METTL6 with single-point mutants in METTL6: K43A, D46A, L47A, K50A, located in METTL6 helix αN2; and R114A, located in the core of METTL6 (Figure 2f). For each mutant, we tested a set of parameters, including binding of METTL6 to tRNA^Ser^ via fluorescence polarization, cofactor binding (SAM, SAH, SAM-competitive inhibitor sinefungin) of METTL6 via a thermal shift assay, formation of the METTL6-SerRS-tRNA^Ser^ complex via a size exclusion co-elution, and methylation activity of the METTL6-SerRS-tRNA^Ser^ mutant complex via an aptamer-based time-resolved fluorescence resonance energy transfer (TR-FRET) assay that measures the formation of the reaction product SAH (Figure 3, Fig. S13). All our METTL6 variable arm interface mutants interacted with the SAM cofactor or SAM analogues; they all bound tRNA^Ser^ with similar low affinity, demonstrating that the overall structural integrity of METTL6 is maintained (Fig. S13). However, all mutations impaired the formation of the full METTL6-SerRS-tRNA^Ser^ complex in the size exclusion experiment, indicating that interference with the interface between METTL6 and tRNA^Ser^ variable arm compromises association of METTL6 with the SerRS-tRNA^Ser^ complex (Figure 3a). In accordance with defective complex formation, the K43A, D46A, L47A and R114A mutants were inactive in the methylation assay, and the K50A mutant had only very low residual activity (Figures 3b, c). To conclude, our complex formation and methylation activity assays align well with the computational results, indicating the crucial role of the N-terminal helix ɑN2 as an important element for allosteric communication within the complex (Figure 3d).

**Figure 3.**
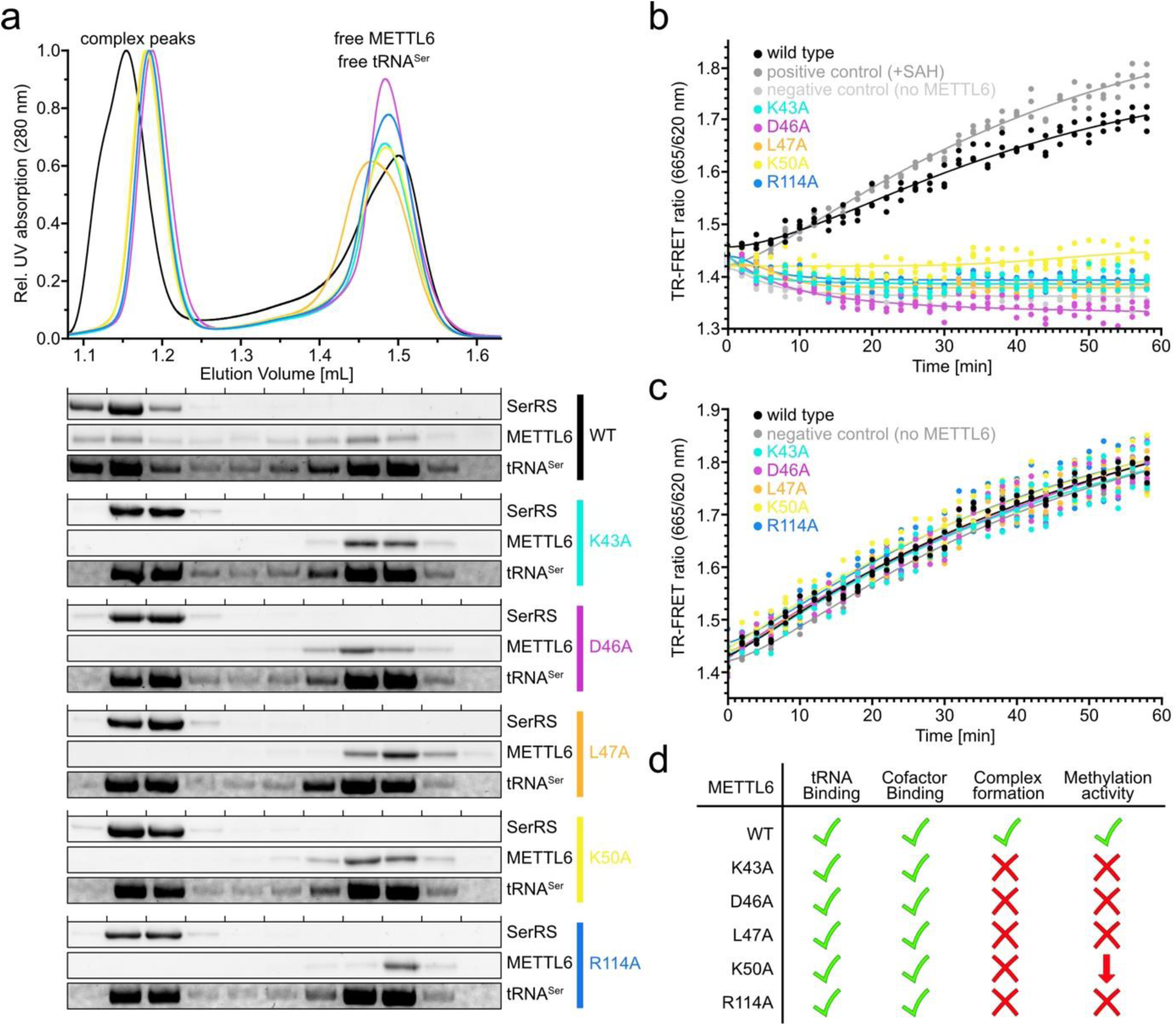
Complex formation and activity of the METTL6-SerRS-tRNA^Ser^ complex. (a) *In vitro* complex assembly of METTL6 mutants with SerRS and tRNA^Ser^ on a Superdex 200 3.2/300 (Cytiva) size-exclusion chromatography (SEC) column. Chromatogram for relative UV absorption at 280 nm is indicated with SDS-PAGE (SerRS, METTL6) and Urea-PAGE (tRNA^Ser^) of the fractions below. (b) *In vitro* methyltransferase activity of different METTL6 variants in the presence of SerRS and tRNA^Ser^. TR-FRET ratios of 665/620 nm emission are indicated. A positive control, including 250 nM SAH instead of SAM and without METTL6, and a negative control containing all reactants but without METTL6 are shown. Non-linear fits of three replicates are shown, with each replicate being indicated as an individual datapoint. (c) Control assay to (b) testing all METTL6 variants in the presence of 250 nM SAH instead of SAM, to exclude direct interference with aptamer formation. TR-FRET ratios of 665/620 nm emissions are indicated. Non-linear fits of three replicates are shown with each replicate being indicated as an individual datapoint. (d) Table summarizing the results of the biochemical experiments.

### Molecular mechanism of tRNA conformational remodeling leading to the flip-out of base C_32_

The conformation of the METTL6-bound tRNA^Ser^ anticodon stem loop is highly distorted in comparison to free tRNA^Ser^ structures. In free tRNA^Ser^, C_32_ is present in a flipped-in conformation, forming a stable non-canonical base pair with A_38_ (Figure 4a) ^70^. Similarly, such C_32_:A^38^ base pairing was also observed in our cryo-EM structure of the METTL6-SerRS-tRNA^Ser^ complex, for the METTL6-free tRNA^Ser^ molecule ^54^. Our MD simulations suggest that SerRS is crucial for correct remodeling of the tRNA^Ser^ anticodon stem loop and for stabilizing the reactive flipped-out conformation of C_32_ that is observed in the METTL6-SerRS-tRNA^Ser^ complex (Figure 4a). To further investigate the molecular mechanism and energetics of this conformational transition in the full METTL6-SerRS-tRNA^Ser^ complex, we performed well-tempered metadynamics simulations ^71^. As collective variables (CVs), we employed eRMSD ^72^ that measures the similarity between two structures by considering only the relative positions and orientations of selected nucleobases (C_32_ and A_38_) and the distance between reacting atoms C_32_:N3 and SAM:CH^3^. This enabled us to sample both the flipped-in and flipped-out conformation (Fig. S14). We determined that in the full complex, the flipped-in and flipped-out conformation were comparable in their relative free energy (ΔG =-0.1 ± 0.6 kcal/mol, Figure 4b). In the flipped-in conformation the C_32_ nucleobase was primarily stabilized by hydrogen bonding to A^38^ and base stacking with A^31^ and U^33^, while C_32_ ribose-phosphate backbone interacted predominantly with Lys58, Arg60, Tyr190 and Arg259 (Figure 4c). Conversely, in the flipped-out conformation, C_32_ nucleobase was held in place by Arg60, Phe158 and Tyr190, with Trp62 and Arg259 stabilizing the backbone (Figure 4c), mirroring the results of our classical MD simulations. Interestingly, the flipped-in and flipped-out state were separated by an intermediate conformation, where the C_32_ is still flipped-in but not engaged in base pairing with A^38^ (Figure 4b). The energetic cost of breaking the non-canonical base pair C_32_:A^38^ was approximately 0.9 ± 0.4 kcal/mol, while the activation free energy barrier (ΔG^ǂ^) was 5.4 ± 0.6 kcal/mol. Instead, the transition from the intermediate to the final flipped-out conformation, where C_32_ is appropriately docked for the methylation reaction to occur, was associated with a lower ΔG^ǂ^ of 2.6 ± 0.5 kcal/mol. This suggests breaking the hydrogen bonds between C_32_ and A^38^ is the rate-limiting step in anticodon stem loop remodeling. In summary, in the full METTL6-SerRS-tRNA^Ser^ complex, where METTL6 is stably docked at the tRNA^Ser^ anticodon stem loop, tRNA^Ser^ conformational remodeling is a low-barrier process, resulting in rapid formation of the reactive flipped-out state.

**Figure 4.**
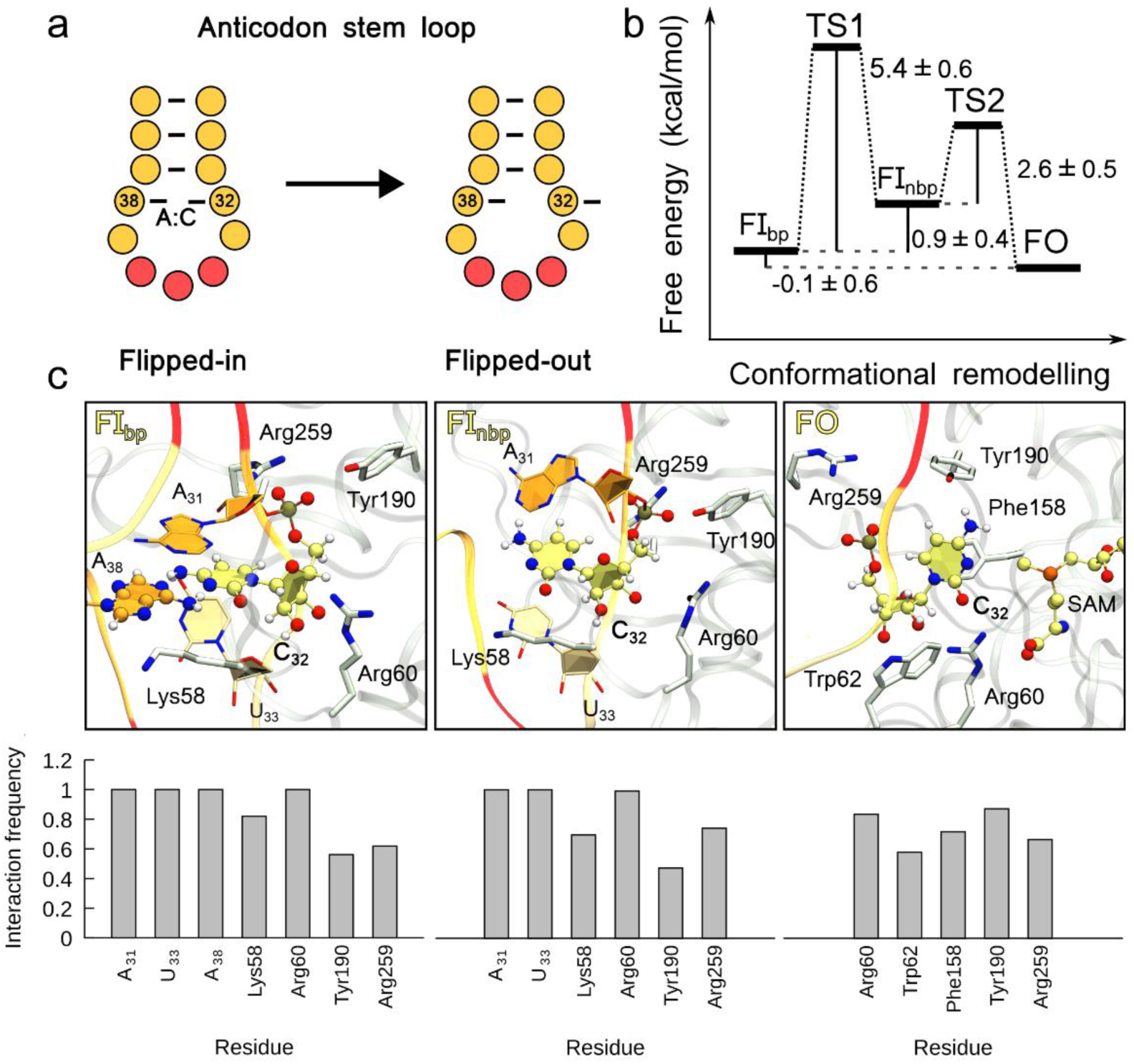
METTL6-SerRS-tRNA^Ser^ complex formation facilitates tRNA^Ser^ conformational remodeling. (a) Scheme of the flipped-in state observed in free tRNA^Ser^ and the flipped-out state observed in METTL6-SerRS-tRNA^Ser^. (b) Free energy profile of the tRNA^Ser^ anticodon stem loop conformational transition from the base paired flipped-in (FI^bp^) to the flipped-out state (FO) as obtained from metadynamics simulation. During anticodon stem loop remodeling a higher-energy intermediate conformation (FI^nbp^) was observed where C_32_ is still flipped-in, but does not form a non-canonical base pair with A^38^. (c) Snapshots of the flipped-in base paired (FI^bp^), flipped-in non-base paired (FI^nbp^) and flipped-out (FO) tRNA^Ser^ anticodon stem loop conformations observed during metadynamics simulations. Residues contributing to the stabilization of C_32_ in the different tRNA^Ser^ conformational states are shown as licorice. For each conformation, interaction persistence of nucleic and protein residues stabilizing C_32_ positioning is shown below.

### Methyl group transfer is a single-step S^N2^-like reaction

In the absence of any obvious acidic or other potentially reactive residues in the proximity of C_32_ or the methyl group of SAM, the catalytic mechanism of the m^3^C formation was unclear.

Chemical reactions cannot be simulated with classical MD, since these cannot capture any phenomena involving electron density transfer, such as bond breaking and formation ^73^. Therefore, we turned to quantum-classical (QM/MM) MD to resolve the precise molecular mechanism of the m^3^C methylation reaction. We partitioned the system into two sections: the quantum (QM) zone, covering the active site (see Methods), and the classical (MM) zone that described the remainder of the system. This allows for a more accurate treatment of the active site and investigation of reaction mechanisms at an affordable computational cost.

To achieve appropriate sampling, we coupled QM/MM MD simulations with umbrella sampling ^74^. The reaction coordinate (RC) was defined as the difference between the length of the breaking bond (SAM:CH^3^-SAM:S) and forming bond (C_32_:N3-SAM:CH^3^) (Figure 5a). Preliminary steered QM/MM MD simulations confirmed this was an appropriate RC for simulating m^3^C_32_ formation (Fig. S15). The reaction was sampled along 16 windows 0.2 Å apart, spanning the RC range from-1.5 Å to 1.5 Å. Each window was sampled 5-10 ps (Fig. S16). Weighted histogram analysis was used to recreate the free energy profile. Methylation of C_32_ to produce m^3^C_32_ occurred in a single step. Our data revealed that the m^3^C methylation reaction followed an S^N2^-like substitution mechanism. The heterocyclic nitrogen atom of C_32_ performed a nucleophilic attack on the electrophilic methyl group of SAM, activated by the presence of the positively charged sulfonium cation. First, the carbon–sulphur bond was cleaved, followed by formation of the nitrogen–carbon bond (Figure 5a). The transition state (TS) was observed at positive values of RC, indicating a late TS with the leaving group departure more advanced than the nucleophilic attack (Figures 5a-c). Indeed, in the TS, the methyl group cation is in its unbound state, surrounded by H^2^O molecules. The uncharged S-adenosyl-L-homocysteine (SAH) was released as the leaving group, and the resulting methylated nucleoside carried a positive charge, with no m^3^C_32_ deprotonation event observed in the product state. The observed activation free energy barrier was 16.6 ± 0.5 kcal/mol, a value characteristic for enzyme catalysis ^75^, while the reaction free energy was-15.4 ± 0.4 kcal/mol, in line with the extremely favorable energetics (free energy change of −17 kcal/mol) for conversion of SAM to SAH (Figure 5b). Taken together, our simulations suggest that METTL6 primarily facilitates the spontaneous methylation reaction by the appropriate geometric positioning of reactant species. The catalytic pocket of m^3^C-tRNA-MTases is highly conserved (Fig. S17) ^41^. Indeed, C_32_ and SAM binding residues are almost identical among human m^3^C-tRNA-MTases (Fig. S17). This indicates the uncovered catalytic mechanism is likely shared across family members.

**Figure 5.**
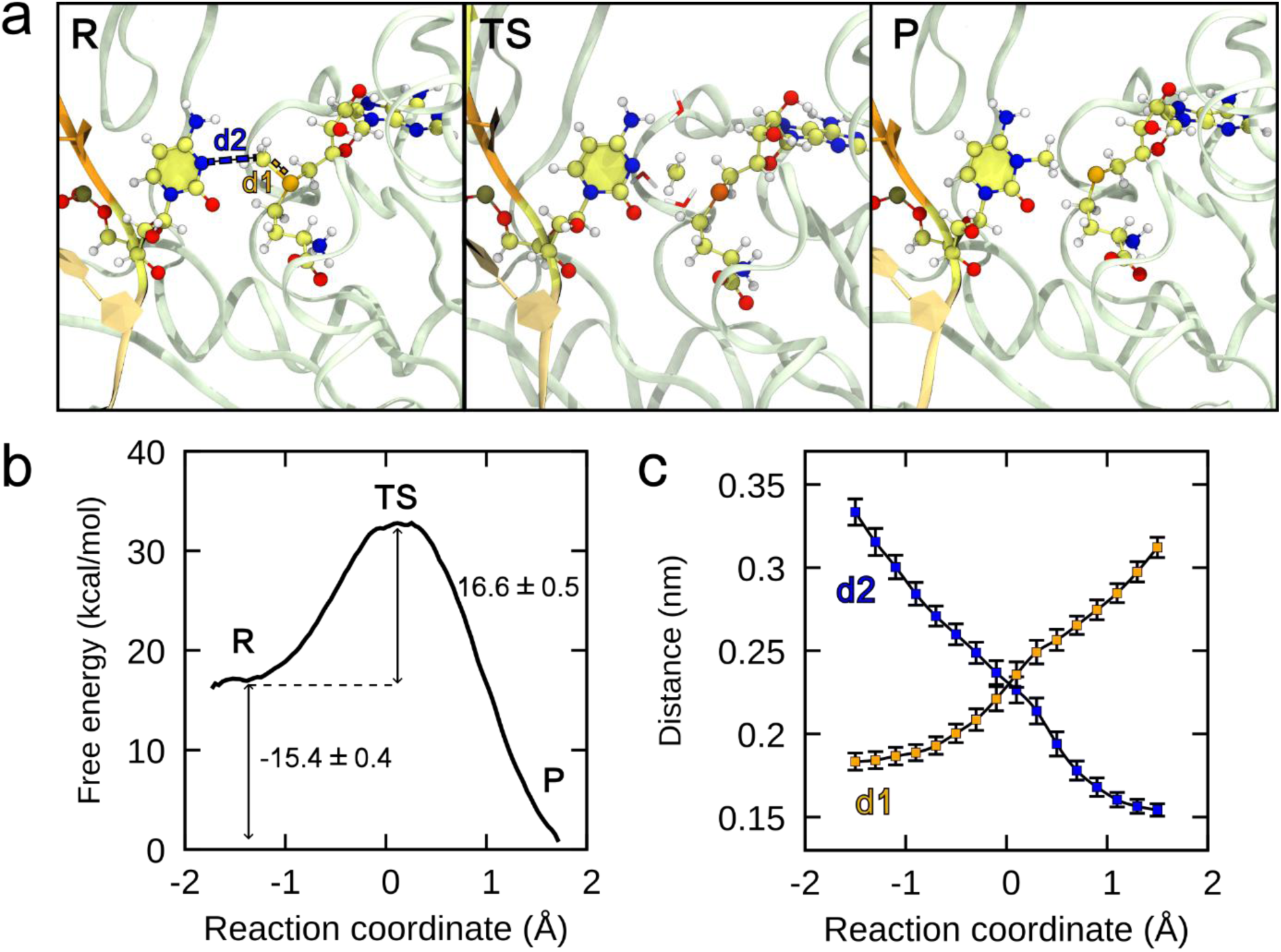
m^3^C methylation is a concerted S^N2^ reaction. (a) Snapshots of the active site in the reactant (R), transition (TS) and product state (P). Distances used for the definition of the reaction coordinate are shown in orange (d1, SAM:CH^3^-SAM:S) and blue (d2, C_32_:N3-SAM:CH^3^). (b) Helmholtz free energy as a function of the reaction coordinate. The reactant (R) and the product state (P) are separated by a single transition state (TS). (c) Values of distances d1 and d2 as defined in panel (a) as a function of the reaction coordinate. Error bars denote ± s.d. from the mean for *n* ≥ 12000 simulation frames.

## Discussion

Through computational methods combined with a panel of *in vitro* biochemical data, we reveal the catalytic mechanism of the m^3^C-tRNA-MTase METTL6. We determine an S^N^2 reaction mechanism for the m^3^C formation. The reaction occurs spontaneously once tRNA^Ser^ and SAM are positioned appropriately within the METTL6-SerRS-tRNA^Ser^ complex, without the aid of any other catalytic residue. Importantly, our computer simulations revealed that SerRS directs the formation of the catalytically favorable geometry and subsequent deposition of m^3^C_32_ through stabilizing the tRNA^Ser^ long variable arm, which leads to onboarding of METTL6 foremost through its N-terminal helices.

The sequences of the N-terminal helices in the m^3^C-tRNA-MTase family members METTL2, METTL6 and METTL8 differ, indicating their central role for tRNA substrate binding specificity, in line with the results of our simulations. METTL2 and METTL8 contain additional conserved sequences in their N-termini - absent in METTL6 - suggesting further alternative regulation elements in this region. Notably, the deletion of this METTL2 and METTL8 exclusive motif in isoform4 of METTL8 abolished its methylating activity ^76^; the same was true for mutants in conserved residues in yeast homologues ^76^. However, the N-terminal helical motif is not the sole determinant for tRNA substrate specificity as the N-terminal domain-swapped chimeras of METTL6 and METTL2 enzymes did not switch their substrate preference and were inactive ^48^.

All m^3^C-tRNA-MTases can associate with protein cofactors. In case of METTL2 and METTL8, their regulatory mechanisms and functional importance for tRNA substrate recognition and methylation remain ambiguous ^49,51,76–78^. On the other hand, the strict requirement of a protein cofactor for efficient methylating activity has been observed for several other RNA-MTases ^37,77^. These cofactors exhibit diverse mechanisms of methyltransferase activation and can function by promoting binding of the methylating ligand and/or substrate tRNA. Examples of such RNA MTase-protein cofactor complexes include METTL3-METTL14 ^79^, where METTL3 is the catalytic domain and METTL14 serves as the inactive scaffold with degenerate active site residues ^80,81^, METTL1-WDR4 ^82,83^ and FTSJ1-THADA ^84^. While some cofactors, like METTL14, function solely as facilitators of RNA modification, others, like SerRS, have also other cellular roles. For example, WDR4, which activates METTL1, also aids genome stability by regulating activity of FEN1 endonuclease ^85^. How often aminoacyl-tRNA synthetases are involved in the activation of tRNA-MTases is unclear and besides SerRS hitherto no other example has been reported in the literature. Interestingly, METTL1 was recently reported to associate with multi-tRNA synthetase complex, promoting tRNA aminoacylation independent of its methylating activity ^86^. Further, METTL8 was found to promote aminoacylation activity in mitochondrial mtSerRS (also termed SerRS2) and mitochondrial threonyl-tRNA synthetase mtTARS (also termed TARS2) ^76^. These findings point to a probable regulatory two-way aminoacyl-tRNA synthetase-methyltransferase axis.

The here uncovered catalytic mechanism of m^3^C formation is likely shared across family members. Moreover, this mechanism is highly reminiscent of that observed in the m^3^C DNA methyltransferase DNMT3A ^87,88^. There, static quantum chemistry calculations, performed using minimal models of the active site in implicit solvent, indicated that the methyl group transfer from SAM to the N3 atom of the target cytidine occurs directly and without any subsequent deprotonation of the positively charged m^3^C nucleotide ^87^. Additionally, structural and mutational studies pointed to the importance of a surrounding Arg792 residue in DNMT3A, whose position overlaps with that of Arg60 in METTL6, for the positioning of the reactive cytidine ^88^. Interestingly in DNMT3A, the positive charge of m^3^C appeared to be stabilized through hydrogen bonding between the m^3^C amino group and a proximal Glu756 residue, which has no analog in METTL6 ^88^. Among the handful of RNA-MTases, whose mode of action has been clarified through structural and computational studies, the catalytic mechanism of METTL6 most resembles that of methyltransferases targeting other nucleophilic nitrogen heteroatoms or exocyclic amino groups. Among these are METTL1 and METTL3 that catalyze the formation of m^7^G and m^6^A, respectively ^37,83,89^. Specifically, QM/MM metadynamics simulations based on the crystal structure of the METTL3-METTL14 complex revealed a direct transfer of the methyl cation from SAM to the N6 atom of the target adenosine prior to any deprotonation event. The reaction occurred by overcoming an activation free energy barrier of 15-16 kcal/mol ^80,89^. However, the authors speculated that the positive charge of the resulting m^6^A could be relieved through deprotonation of the m^6^NH^2+^ group by surrounding water molecules after the methylation reaction. Most importantly, their results suggest the residues lining the catalytic cavity appear to primarily ensure the correct positioning of the reactive species without directly participating in the methylation reaction, which aligns with what we observed here for METTL6. Similarly, the structure of the recently resolved METTL1-WDR4 complex suggests m^7^G formation also occurs as a direct S^N^2-like reaction ^83^. Interestingly, the catalytic cavity of METTL1 contains several negatively charged residues (Asp163, Asp199 and Glu240) whose mutation to Ala was detrimental to catalytic activity. However, as no computational study has been performed, it is unclear whether these serve to provide a favorable electrostatic environment that stabilizes the methyl cation in the transition state and/or the positively charged m^7^G residue in the product state, or, instead, mediate m^7^G deprotonation at the N1 position, dissipating the positive charge. Conversely, formation of m^5^C, catalyzed be the NOL1/NOP2/Sun family of RNA-MTases, occurs through a completely distinct reaction pathway, involving two highly conserved cysteine residues, multiple bond rearrangements and a covalent protein-RNA intermediate ^90^.

## Conclusions

In conclusion, our work firmly establishes the catalytic and recognition mechanisms of METTL6 in molecular detail. While the catalytic mechanism of m^3^C formation appears broadly shared among m^3^C-tRNA-MTases, future experimental structural investigation will reveal fine details of their differential substrate-selection mechanisms. Further, additional detailed investigations will be necessary to reveal the precise functional role of other RNA modifications in the anticodon stem loop (i^6^A^37^, t^6^A^37^) on C_32_ methylation. Finally, our outcomes provide concrete indications for viable therapeutic approaches aimed at inhibiting tRNA^Ser^ C_32_ methylation. According to our analysis of the METTL6-tRNA^Ser^ variable arm interface, several non-redundant, albeit essential residues render the interaction fragile and could thus potentially provide an Achille’s heel for therapeutic targeting of METTL6. The apparent specificity of the SerRS-aided METTL6 recognition mechanism suggests that focusing on the METTL6-SerRS-tRNA^Ser^ tertiary interface may lead to the discovery of METTL6-specific therapeutics. In contrast, targeting the active site in m^3^C-tRNA-MTases could lead to the development of multi-specificity drugs that modulate the entire m^3^C metabolism, offering broader therapeutic effects, but also potential off-target effects. Due to the interplay of m^3^C-tRNA-MTases observed in different cancer types, their poor selectivity may be particularly beneficial and result in synergistic effects ^40,60,91^.

## Materials and Methods

### Classical molecular dynamics (MD)

As the starting point for our molecular dynamics (MD) simulations, we utilized the 1:2:2 METTL6-SerRS-tRNA^Ser^ complex resolved with cryo-EM (PDBID: 8P7B) ^54^. To construct the smaller bimolecular METTL6-tRNA^Ser^ complex, SerRS molecules and the METTL6-unbound tRNA^Ser^ were removed from the complex. System topologies were prepared with the CHARMM-GUI web platform ^92^. As force field, we employed CHARMM36m that contains parametrizations for all present modified RNA nucleotides ^65,93^. Systems were placed in a cubic box and solvated with TIP3P water ^94^. KCl salt was added to 150 mM concentration. For simulations performed with the Amber force field, we utilized ff14SB and χ^OL3^ force fields for protein and RNA, respectively. RNA modifications were modelled with the modXNA tool ^95^. Parameters for SAM were obtained with the antechamber tool, using amber atom types and AM1-BCC charges ^96^. Ions were described with Li-Merz parameters compatible with the TIP3P water model ^97^. Classical all-atom MD simulations were performed in GROMACS 2022.3 ^66^. Each simulated system was first equilibrated in the NVT ensemble for 10 ns using periodic boundary conditions and position restraints on the protein and RNA components. Next, equilibration was continued in the NPT ensemble for another 10 ns. Afterwards, position restraints were removed and NPT equilibration was extended for another 10 ns. Newton’s equations of motion were solved with the leap-frog algorithm using a 1 fs time step. Electrostatic interactions were evaluated using the particle mesh Ewald method ^98^. Temperature (T) was controlled using the V-rescale thermostat (T = 300 K, coupling constant was 0.1 ps), while pressure was kept at 1 bar with the Parrinello-Rahman barostat (time constant for pressure coupling was 2 ps) ^99,100^. Bonds involving hydrogen atoms were constrained with the LINCS algorithm. For van der Waals interaction, force-switching was applied with 1 nm cutoff. Cutoff for electrostatic and van der Waals interactions was set at 1.2 nm. Production runs were performed in triplicates, each run 500 ns in length, to obtain reliable statistics. Trajectory analysis was performed with the python package MDAnalysis ^101^. Interactions were analyzed with the python package prolif ^102^. Dynamical network analysis was performed with the python package dynetan ^67^. Dynamical network analysis was carried out on the MD trajectory of the METTL6-SerRS-tRNA^Ser^ complex obtained by concatenating the three replica simulations (totaling 1.5 μs). Each residue was represented by one node. For protein and nucleic residues, Cα and P atoms were selected as representative atoms for node construction, respectively. For SAM, the sulphur atom represented the node. For each simulation frame, a residue-residue contact map was calculated utilizing the cutoff distance of 4.5 Å and considering only the positions of heavy atoms. Two nodes were considered in direct communication if their contact persistency was above 60 %. Optimal paths were calculated using the Floyd Warshall algorithm. These were then used to determine edge betweenness. The network was then partitioned into communities using Louvain heuristics. To obtain a coarse-grained betweenness value corresponding to the communication strength between two selected communities, we summed edge betweenness values of all residue pairs connecting the selected communities.

### Metadynamics simulations

Conformational remodeling of the tRNA^Ser^ anticodon stem loop in the full METTL6-SerRS-tRNA^Ser^ complex was investigated with well-tempered metadynamics ^103^. Metadynamics simulations were performed in GROMACS 2022.3 patched with PLUMED 2.8.1 ^72^. As collective variables we employed eRMSD between the nucleotide pair C_32_:A^38^ and the distance between reacting atoms C_32_:N3 and SAM:CH^3^. Simulations were performed at 300 K. Bias factor was set at 15 and the tau parameter, which determines the rate of Gaussian hill height damping, at 70. The width of the Gaussian hills was 0.1 Å. A Gaussian hill was added to the potential energy surface every 250 steps. We performed a metadynamics run (∼ 1.5 μs) to map out the full 2D free energy surface in order to determine all free energy minima and the low-barrier pathways connecting them. We identified three minima, corresponding to the flipped-in base pairing (FI^bp^) conformation, the flipped-in non-base pairing (FI^nbp^) conformation and the flipped-out (FO) conformation as described in the main text. The free energy profile was calculated with the sum_hills utility provided with PLUMED ^72^. Error analysis was performed utilizing the block averaging method.

### Hybrid quantum-classical (QM/MM) molecular dynamics (MD)

QM/MM MD simulations were performed with GROMACS 2023.3 utilizing its cp2k interface (version 2024.1) ^104^. The QM portion was composed of the target nucleotide C_32_, methyl donor SAM and surrounding water molecules. The quantum box therefore contained 78 atoms. The QM portion was described with the Becke-Lee-Yang-Parr density functional theory (DFT/BLYP) ^105,106^. Double-ζ molecularly optimized basis set was used along with Goedecker– Teter–Hutter pseudopotentials (DZVP-MOLOPT-GTH) ^107^. Plane wave cutoff was set at 400 Ry and DFT-D3 dispersion correction was applied ^108^. First, the system was relaxed in the NVT ensemble for 5 ps. MD run parameters equaled those applied during classical MD, however the time step was reduced to 0.5 fs. Following system equilibration, umbrella sampling simulations were performed to obtain the reaction free energy profile ^74,109^. We employed a single reaction coordinate (RC), defined as the difference in length of the breaking bond (SAM:CH^3^ – SAM:S) and forming bond (C_32_:N3 – SAM:CH^3^). We scanned the RC in the-1.5 Å to 1.5 Å range using a 0.2 Å step size, resulting in 16 simulation windows. The harmonic force constant was set at 750 kJ mol^-1^ Å^-2^, to achieve a good overlap between RC distributions of neighboring sampling windows. At each RC value, the system was simulated for 5-10 ps. Free energy profile was reconstructed from pull forces obtained for each simulation window using the weighted histogram analysis method (WHAM) utilizing the gmx wham tool ^110^. Errors were calculated with the bootstrapping method utilizing 200 bootstraps.

### Protein expression and purification

SerRS and METTL6 variants were expressed and purified as previously described ^54^. Concentrated samples were flash-frozen in liquid nitrogen and stored at-80 °C.

### In vitro transcription of tRNA^Ser^

Human tRNA^Ser^ was in vitro transcribed from a pUC19 vector using T7 polymerase runoff transcription as previously described ^54^.

### Fluorescence polarization assay

In vitro transcribed tRNA^SerUGA^ was 3’-Fluorescein labeled by mixing 100 µg tRNA 1:1 with a 2X oxidation solution containing 0.2 M sodium periodate and 0.2 M sodium acetate. The mixture was incubated at room temperature for 1.5 h. The oxidation reaction was quenched by the addition of KCl to a final concentration of 0.25 M. Resulting KIO^4^ was removed by centrifugation. Fluorescein-5-thiosemicarbazide was added to the supernatant to a final concentration of 50 mM and incubated overnight in the dark at room temperature. Labeled tRNAs were urea-PAGE purified and refolded. Serial dilutions of each METTL6 variant starting at final concentrations of 150 µM were each incubated with fluorescein-labeled tRNA^SerUGA^ at a final concentration of 25 nM for 15 min at room temperature in a buffer containing 20 mM HEPES-KOH (pH 7.5), 50 mM NaCl, 2 mM MgCl^2^, 1mM DTT, 0.1 % TWEEN-20, 0.5 U µl^-1^ RNasin Plus RNase Inhibitor (Promega) and 0.5 mM sinefungin (Merck). Fluorescence polarization was measured with a ClarioStar (BMG Labtech) plate reader in 384-well small volume flat black microplates (Greiner) with excitation/emission wavelengths at 460/515 nm. Fluorescein emissions were detected at 515 nm. Data was fitted using the sigmoidal fit interpolation (GraphPad Prism).

### Thermal shift assay

METTL6 variants were diluted to final concentrations of 20 µM each in a buffer containing 20 mM HEPES-KOH (pH 7.5), 100 mM NaCl, 2 mM MgCl^2^, 1 mM DTT and 2x SYPRO Orange (Invitrogen). The METTL6 cofactors SAM, SAH and sinefungin were individually added to final concentrations of 1 mM each. The METTL6 variants in the presence or absence of different cofactors were exposed to a temperature gradient ranging from 10 °C to 100 °C with an increase of 1 °C each minute. The fluorescence was measured using real-time PCR (CFX96, Biorad). The minimum/maximum value ratio for a temperature range from 35 °C to 75 °C were plotted in GraphPad prism using the normalization function.

### Size exclusion chromatography (SEC) complex assembly assay

The complex assembly of METTL6 variants with SerRS and tRNA^Ser^ was investigated with analytical SEC using a Superdex 200 3.2/300 column (Cytiva). METTL6, SerRS and tRNA^Ser^ (at 30 µM each) were incubated on ice for 10 min together with 1 mM sinefungin. Formed complexes were loaded onto the pre-equilibrated SEC column and complexes were eluted in complex formation buffer containing 20 mM HEPES-KOH (pH 7.5), 50 mM NaCl, 2 mM MgCl^2^ and 0.5 mM TCEP. Protein content of each fraction was determined by SDS-PAGE stained with Coomassie blue and RNA content by urea-PAGE stained with methylene blue.

### Methyltransferase activity assay

SAH release during methyltransferase reaction was detected using an artificial split-aptamer derived from the naturally occurring SAH-binding metH aptamer ^111–113^ and the signal was followed using time-resolved Förster resonance energy transfer (TR-FRET). Activity assays were performed in 20 µL of a buffer containing 10 mM HEPES-KOH (pH 7.9), 50 mM KCl, 1.5 mM MgCl^2^, 1 mM DTT and 0.01% TWEEN-20 with 250 nM METTL6 variants and 500 nM SerRS and tRNA^Ser^. The aptamer halves P1 and P2 (see SI) were added to final concentrations of 15 nM and 30 nM, respectively, and Eu^3+^-Streptavidin was added to 0.5 nM. To start the reaction, SAM was added to a final concentration of 2 µM. All assays were performed at room temperature in 384-well small volume flat black microplates (Greiner) and measured using a EnVision 2104 multimode plate reader (Perkin Elmer). The TR-FRET signal originating from metH-SAH complex formation was determined by exciting the donor at 320 nm and measuring the emission at 620 nm and 665 nm, respectively. The TR-FRET ratios of 665/620 nm emission at any given time point were fitted in GraphPad Prism using a non-linear fit (GraphPad).

## ASSOCIATED CONTENT

### Supporting Information

The following files are available.

Supporting Figures S1-S17 depicting additional results obtained by analysis of molecular dynamics trajectories and biochemical characterization of METTL6 mutants’ interactions with tRNA^Ser^ and SAM analogues; Supporting Table S1 listing employed aptamer sequences. (PDF)

## AUTHOR INFORMATION

### Corresponding Author

Eva Kowalinski - European Molecular Biology Laboratory, Grenoble, France;

Alessandra Magistrato - CNR - Istituto Officina dei Materiali (IOM) at International School for Advanced Studies (SISSA), Trieste, Italy;

### Notes

The authors declare no competing financial interest.

## Acknowledgments

We acknowledge the CINECA award under the ISCRA initiative, for the availability of high performance computing resources and support. The authors acknowledge Martin Pelosse for support in using the Eukaryotic Expression Facility at EMBL Grenoble. We thank the HTX Team at EMBL Grenoble for assistance with the thermostability assay. This work used the platforms of the Grenoble Instruct-ERIC Centre (ISBG; UAR 3518 CNRS-CEA-UGA-EMBL) within the Grenoble Partnership for Structural Biology (PSB), supported by FRISBI (ANR-10-INBS-0005-02) and GRAL, financed within the University Grenoble Alpes graduate school (Écoles Universitaires de Recherche) CBH-EUR-GS (ANR-17-EURE-0003). We thank the EMBL Chemical Biology Core Facility (CBCF) for useful discussions, support and use of their equipment. J.A. and A.M. thank the PNRR: National Center for Gene Therapy and Drugs based on RNA Technology CUP B83C22002860006 CN_0000004. A.M. is supported by the Italian Association for Cancer research (AIRC IG 24514).

## References

(1) Roundtree, I. A.; Evans, M. E.; Pan, T.; He, C. Dynamic RNA Modifications in Gene Expression Regulation. Cell 2017, 169 (7), 1187–1200. 10.1016/j.cell.2017.05.045.

(2) Gilbert, W. V.; Bell, T. A.; Schaening, C. Messenger RNA Modifications: Form, Distribution, and Function. Science 2016, 352 (6292), 1408–1412. 10.1126/science.aad8711.

(3) Ontiveros, R. J.; Stoute, J.; Liu, K. F. The Chemical Diversity of RNA Modifications. Biochem. J. 2019, 476 (8), 1227–1245. 10.1042/BCJ20180445.

(4) Motorin, Y.; Helm, M. RNA Nucleotide Methylation: 2021 Update. WIREs RNA 2022, 13 (1), e1691. 10.1002/wrna.1691.

(5) Machnicka, M. A.; Olchowik, A.; Grosjean, H.; Bujnicki, J. M. Distribution and Frequencies of Post-Transcriptional Modifications in TRNAs. RNA Biol. 2014, 11 (12), 1619–1629. 10.4161/15476286.2014.992273.

(6) Jackman, J. E.; Alfonzo, J. D. Transfer RNA Modifications: Nature’s Combinatorial Chemistry Playground. Wiley Interdiscip. Rev. RNA 2013, 4 (1), 35–48. 10.1002/WRNA.1144.

(7) Wang, X.; He, C. Dynamic RNA Modifications in Posttranscriptional Regulation. Mol. Cell 2014, 56 (1), 5–12. 10.1016/j.molcel.2014.09.001.

(8) Zhao, B. S.; Roundtree, I. A.; He, C. Post-Transcriptional Gene Regulation by MRNA Modifications. Nat. Rev. Mol. Cell Biol. 2017, 18 (1), 31–42. 10.1038/nrm.2016.132.

(9) Gilbert, W. V.; Nachtergaele, S. MRNA Regulation by RNA Modifications. Annu. Rev. Biochem. 2023, 92 (1), 175–198. 10.1146/annurev-biochem-052521-035949.

(10) Frye, M.; Harada, B. T.; Behm, M.; He, C. RNA Modifications Modulate Gene Expression during Development. Science (80-.). 2018, 361 (6409), 1346–1349. 10.1126/science.aau1646.

(11) Gustilo, E. M.; Vendeix, F. A.; Agris, P. F. TRNA’s Modifications Bring Order to Gene Expression. Curr. Opin. Microbiol. 2008, 11 (2), 134–140. 10.1016/j.mib.2008.02.003.

(12) Duechler, M.; Leszczyńska, G.; Sochacka, E.; Nawrot, B. Nucleoside Modifications in the Regulation of Gene Expression: Focus on TRNA. Cell. Mol. Life Sci. 2016, 73 (16), 3075–3095. 10.1007/s00018-016-2217-y.

(13) Dedon, P. C.; Begley, T. J. Dysfunctional TRNA Reprogramming and Codon-Biased Translation in Cancer. Trends Mol. Med. 2022, 28 (11), 964–978. 10.1016/j.molmed.2022.09.007.

(14) Manickam, N.; Joshi, K.; Bhatt, M. J.; Farabaugh, P. J. Effects of TRNA Modification on Translational Accuracy Depend on Intrinsic Codon-Anticodon Strength. Nucleic Acids Res. 2015, 44 (4), 1871–1881. 10.1093/nar/gkv1506.

(15) Endres, L.; Fasullo, M.; Rose, R. TRNA Modification and Cancer: Potential for Therapeutic Prevention and Intervention. Future Med. Chem. 2019, 11 (8), 885–900. 10.4155/fmc-2018-0404.

(16) Rapino, F.; Delaunay, S.; Rambow, F.; Zhou, Z.; Tharun, L.; De Tullio, P.; Sin, O.; Shostak, K.; Schmitz, S.; Piepers, J.; Ghesquière, B.; Karim, L.; Charloteaux, B.; Jamart, D.; Florin, A.; Lambert, C.; Rorive, A.; Jerusalem, G.; Leucci, E.; Dewaele, M.; Vooijs, M.; Leidel, S. A.; Georges, M.; Voz, M.; Peers, B.; Büttner, R.; Marine, J. C.; Chariot, A.; Close, P. Codon-Specific Translation Reprogramming Promotes Resistance to Targeted Therapy. Nature 2018, 558 (7711), 605–609. 10.1038/s41586-018-0243-7.

(17) Tuorto, F.; Lyko, F. Genome Recoding by TRNA Modifications. Open Biol. 2016, 6 (12), 160287. 10.1098/rsob.160287.

(18) Watkins, C. P.; Zhang, W.; Wylder, A. C.; Katanski, C. D.; Pan, T. A Multiplex Platform for Small RNA Sequencing Elucidates Multifaceted TRNA Stress Response and Translational Regulation. Nat. Commun. 2022, 13 (1), 2491. 10.1038/s41467-022-30261-3.

(19) Nedialkova, D. D.; Leidel, S. A. Optimization of Codon Translation Rates via TRNA Modifications Maintains Proteome Integrity. Cell 2015, 161 (7), 1606–1618. 10.1016/j.cell.2015.05.022.

(20) Grosjean, H.; Westhof, E. An Integrated, Structure-and Energy-Based View of the Genetic Code. Nucleic Acids Res. 2016, 44 (17), 8020–8040. 10.1093/nar/gkw608.

(21) Cui, J.; Sendinc, E.; Liu, Q.; Kim, S.; Fang, J. Y.; Gregory, R. I. M3C32 TRNA Modification Controls Serine Codon-Biased MRNA Translation, Cell Cycle, and DNA-Damage Response. Nat. Commun. 2024, 15 (1), 1–14. 10.1038/s41467-024-50161-y.

(22) Biela, A. D.; Nowak, J. S.; Biela, A. P.; Mukherjee, S.; Moafinejad, S. N.; Maiti, S.; Chramiec-Głąbik, A.; Mehta, R.; Jeżowski, J.; Dobosz, D.; Dahate, P.; Arluison, V.; Wien, F.; Indyka, P.; Rawski, M.; Bujnicki, J. M.; Lin, T. Y.; Glatt, S. Determining the Effects of Pseudouridine Incorporation on Human TRNAs. EMBO J. 2025, 44 (13), 3553–3585. 10.1038/s44318-025-00443-y.

(23) Schultz, S. K.; Katanski, C. D.; Halucha, M.; Peña, N.; Fahlman, R. P.; Pan, T.; Kothe, U. Modifications in the T Arm of TRNA Globally Determine TRNA Maturation, Function, and Cellular Fitness. Proc. Natl. Acad. Sci. U. S. A. 2024, 121 (26), e2401154121. 10.1073/pnas.2401154121.

(24) Rak, R.; Polonsky, M.; Eizenberg-Magar, I.; Mo, Y.; Sakaguchi, Y.; Mizrahi, O.; Nachshon, A.; Reich-Zeliger, S.; Stern-Ginossar, N.; Dahan, O.; Suzuki, T.; Friedman, N.; Pilpel, Y. Dynamic Changes in TRNA Modifications and Abundance during T Cell Activation. Proc. Natl. Acad. Sci. U. S. A. 2021, 118 (42). 10.1073/pnas.2106556118.

(25) Linder, B.; Sharma, P.; Wu, J.; Birbaumer, T.; Eggers, C.; Murakami, S.; Ott, R. E.; Fenzl, K.; Vorgerd, H.; Erhard, F.; Jaffrey, S. R.; Leidel, S. A.; Steinmetz, L. M. TRNA Modifications Tune M6A-Dependent MRNA Decay. Cell 2025, 188 (14), 3715–3727.e13. 10.1016/j.cell.2025.04.013.

(26) Giguère, S.; Wang, X.; Huber, S.; Xu, L.; Warner, J.; Weldon, S. R.; Hu, J.; Phan, Q. A.; Tumang, K.; Prum, T.; Ma, D.; Kirsch, K. H.; Nair, U.; Dedon, P.; Batista, F. D. Antibody Production Relies on the TRNA Inosine Wobble Modification to Meet Biased Codon Demand. Science (80-.). 2024, 383 (6679), 205–211. 10.1126/science.adi1763.

(27) Endres, L.; Dedon, P. C.; Begley, T. J. Codon-Biased Translation Can Be Regulated by Wobble-Base TRNA Modification Systems during Cellular Stress Responses. RNA Biol. 2015, 12 (6), 603–614. 10.1080/15476286.2015.1031947.

(28) Deng, W.; Babu, I. R.; Su, D.; Yin, S.; Begley, T. J.; Dedon, P. C. Trm9-Catalyzed TRNA Modifications Regulate Global Protein Expression by Codon-Biased Translation. PLoS Genet. 2015, 11 (12), e1005706. 10.1371/journal.pgen.1005706.

(29) Guo, H.; Xia, L.; Wang, W.; Xu, W.; Shen, X.; Wu, X.; He, T.; Jiang, X.; Xu, Y.; Zhao, P.; Tan, D.; Zhang, X.; Zhang, Y. Hypoxia Induces Alterations in TRNA Modifications Involved in Translational Control. BMC Biol. 2023, 21 (1), 39. 10.1186/s12915-023-01537-x.

(30) Gu, C.; Begley, T. J.; Dedon, P. C. TRNA Modifications Regulate Translation during Cellular Stress. FEBS Lett. 2014, 588 (23), 4287–4296. 10.1016/j.febslet.2014.09.038.

(31) Rappol, T.; Waldl, M.; Chugunova, A.; Hofacker, I. L.; Pauli, A.; Vilardo, E. TRNA Expression and Modification Landscapes, and Their Dynamics during Zebrafish Embryo Development. Nucleic Acids Res. 2024, 52 (17), 10575–10594. 10.1093/nar/gkae595.

(32) Delaunay, S.; Helm, M.; Frye, M. RNA Modifications in Physiology and Disease: Towards Clinical Applications. Nat. Rev. Genet. 2024, 25 (2), 104–122. 10.1038/s41576-023-00645-2.

(33) Orellana, E. A.; Siegal, E.; Gregory, R. I. TRNA Dysregulation and Disease. Nat. Rev. Genet. 2022, 23 (11), 651–664. 10.1038/s41576-022-00501-9.

(34) Jonkhout, N.; Tran, J.; Smith, M. A.; Schonrock, N.; Mattick, J. S.; Novoa, E. M. The RNA Modification Landscape in Human Disease. RNA 2017, 23 (12), 1754–1769. 10.1261/rna.063503.117.

(35) Wang, L.; Lin, S. Emerging Functions of TRNA Modifications in MRNA Translation and Diseases. J. Genet. Genomics 2023, 50 (4), 223–232. 10.1016/j.jgg.2022.10.002.

(36) Suzuki, T. The Expanding World of TRNA Modifications and Their Disease Relevance. Nat. Rev. Mol. Cell Biol. 2021, 22 (6), 375–392. 10.1038/s41580-021-00342-0.

(37) Höbartner, C.; Bohnsack, K. E.; Bohnsack, M. T. How Natural Enzymes and Synthetic Ribozymes Generate Methylated Nucleotides in RNA. Annu. Rev. Biochem. 2024, 93 (1), 109–137. 10.1146/annurev-biochem-030222-112310.

(38) Martin, J. SAM (Dependent) I AM: The S-Adenosylmethionine-Dependent Methyltransferase Fold. Curr. Opin. Struct. Biol. 2002, 12 (6), 783–793. 10.1016/S0959-440X(02)00391-3.

(39) Oerum, S.; Meynier, V.; Catala, M.; Tisné, C. A Comprehensive Review of M6A/M6Am RNA Methyltransferase Structures. Nucleic Acids Res. 2021, 49 (13), 7239–7255. 10.1093/nar/gkab378.

(40) Ignatova, V. V.; Kaiser, S.; Ho, J. S. Y.; Bing, X.; Stolz, P.; Tan, Y. X.; Lee, C. L.; Gay, F. P. H.; Lastres, P. R.; Gerlini, R.; Rathkolb, B.; Aguilar-Pimentel, A.; Sanz-Moreno, A.; Klein-Rodewald, T.; Calzada-Wack, J.; Ibragimov, E.; Valenta, M.; Lukauskas, S.; Pavesi, A.; Marschall, S.; Leuchtenberger, S.; Fuchs, H.; Gailus-Durner, V.; De Angelis, M. H.; Bultmann, S.; Rando, O. J.; Guccione, E.; Kellner, S. M.; Schneider, R. METTL6 Is a TRNA M3C Methyltransferase That Regulates Pluripotency and Tumor Cell Growth. Sci. Adv. 2020, 6 (35), eaaz4551. 10.1126/sciadv.aaz4551.

(41) Bohnsack, K. E.; Kleiber, N.; Lemus-Diaz, N.; Bohnsack, M. T. Roles and Dynamics of 3-Methylcytidine in Cellular RNAs. Trends Biochem. Sci. 2022, 47 (7), 596–608. 10.1016/j.tibs.2022.03.004.

(42) Berger, K. D.; Puthenpeedikakkal, A. M. K.; Mathews, D. H.; Fu, D. Structural Impact of 3-Methylcytosine Modification on the Anticodon Stem-Loop of a Neuronally-Enriched Arginine TRNA. J. Mol. Biol. 2025, 437 (16), 169096. 10.1016/J.JMB.2025.169096.

(43) Han, L.; Marcus, E.; D’Silva, S.; Phizicky, E. M. S. Cerevisiae Trm140 Has Two Recognition Modes for 3-Methylcytidine Modification of the Anticodon Loop of TRNA Substrates. RNA 2017, 23 (3), 406–419. 10.1261/rna.059667.116.

(44) Noma, A.; Yi, S.; Katoh, T.; Takai, Y.; Suzuki, T.; Suzuki, T. Actin-Binding Protein ABP140 Is a Methyltransferase for 3-Methylcytidine at Position 32 of TRNAs in Saccharomyces Cerevisiae. RNA 2011, 17 (6), 1111–1119. 10.1261/rna.2653411.

(45) D’Silva, S.; Haider, S. J.; Phizicky, E. M. A Domain of the Actin Binding Protein Abp140 Is the Yeast Methyltransferase Responsible for 3-Methylcytidine Modification in the TRNA Anti-Codon Loop. RNA 2011, 17 (6), 1100–1110. 10.1261/rna.2652611.

(46) Arimbasseri, A. G.; Iben, J.; Wei, F. Y.; Rijal, K.; Tomizawa, K.; Hafner, M.; Maraia, R. J. Evolving Specificity of TRNA 3-Methyl-Cytidine-32 (M3C32) Modification: A Subset of TRNAsSer Requires N6-Isopentenylation of A37. RNA 2016, 22 (9), 1400–1410. 10.1261/rna.056259.116.

(47) Xu, L.; Liu, X.; Sheng, N.; Oo, K. S.; Liang, J.; Chionh, Y. H.; Xu, J.; Ye, F.; Gao, Y. G.; Dedon, P. C.; Fu, X. Y. Three Distinct 3-Methylcytidine (M3C) Methyltransferases Modify TRNA and MRNA in Mice and Humans. J. Biol. Chem. 2017, 292 (35), 14695– 14703. 10.1074/jbc.M117.798298.

(48) Mao, X. L.; Li, Z. H.; Huang, M. H.; Wang, J. T.; Zhou, J. B.; Li, Q. R.; Xu, H.; Wang, X. J.; Zhou, X. L. Mutually Exclusive Substrate Selection Strategy by Human M3C RNA Transferases METTL2A and METTL6. Nucleic Acids Res. 2021, 49 (14), 8309–8323. 10.1093/nar/gkab603.

(49) Kleiber, N.; Lemus-Diaz, N.; Stiller, C.; Heinrichs, M.; Mai, M. M. Q.; Hackert, P.; Richter-Dennerlein, R.; Höbartner, C.; Bohnsack, K. E.; Bohnsack, M. T. The RNA Methyltransferase METTL8 Installs M3C32 in Mitochondrial TRNAsThr/Ser(UCN) to Optimise TRNA Structure and Mitochondrial Translation. Nat. Commun. 2022, 13 (1), 1–19. 10.1038/s41467-021-27905-1.

(50) Schöller, E.; Marks, J.; Marchand, V.; Bruckmann, A.; Powell, C. A.; Reichold, M.; Mutti, C. D.; Dettmer, K.; Feederle, R.; Hüttelmaier, S.; Helm, M.; Oefner, P.; Minczuk, M.; Motorin, Y.; Hafner, M.; Meister, G. Balancing of Mitochondrial Translation through METTL8-Mediated M3C Modification of Mitochondrial TRNAs. Mol. Cell 2021, 81 (23), 4810–4825.e12. 10.1016/j.molcel.2021.10.018.

(51) Huang, M. H.; Peng, G. X.; Mao, X. L.; Wang, J. T.; Zhou, J. B.; Zhang, J. H.; Chen, M.; Wang, E. D.; Zhou, X. L. Molecular Basis for Human Mitochondrial TRNA M3C Modification by Alternatively Spliced METTL8. Nucleic Acids Res. 2022, 50 (7), 4012–4028. 10.1093/NAR/GKAC184.

(52) Chen, R.; Zhou, J.; Liu, L.; Mao, X. L.; Zhou, X.; Xie, W. Crystal Structure of Human METTL6, the M3C Methyltransferase. *Commun*. Biol. 2021, 4 (1), 1–10. 10.1038/s42003-021-02890-9.

(53) Li, S.; Zhou, H.; Liao, S.; Wang, X.; Zhu, Z.; Zhang, J.; Xu, C. Structural Basis for METTL6-Mediated M3C RNA Methylation. Biochem. Biophys. Res. Commun. 2022, 589, 159–164. 10.1016/j.bbrc.2021.12.013.

(54) Throll, P.; G. Dolce, L.; Rico-Lastres, P.; Arnold, K.; Tengo, L.; Basu, S.; Kaiser, S.; Schneider, R.; Kowalinski, E. Structural Basis of TRNA Recognition by the M3C RNA Methyltransferase METTL6 in Complex with SerRS Seryl-TRNA Synthetase. Nat. Struct. Mol. Biol. 2024, 31 (10), 1614–1624. 10.1038/s41594-024-01341-3.

(55) Xu, X.; Shi, Y.; Yang, X.-L. Crystal Structure of Human Seryl-TRNA Synthetase and Ser-SA Complex Reveals a Molecular Lever Specific to Higher Eukaryotes. Structure 2013, 21 (11), 2078–2086. 10.1016/j.str.2013.08.021.

(56) Härtlein, M.; Cusack, S. Structure, Function and Evolution of Seryl-TRNA Synthetases: Implications for the Evolution of Aminoacyl-TRNA Synthetases and the Genetic Code. J. Mol. Evol. 1995, 40 (5), 519–530. 10.1007/BF00166620.

(57) Belrhali, H.; Yaremchuk, A.; Tukalo, M.; Berthet-Colominas, C.; Rasmussen, B.; Bösecke, P.; Diat, O.; Cusack, S. The Structural Basis for Seryl-Adenylate and Ap4A Synthesis by Seryl-TRNA Synthetase. Structure 1995, 3 (4), 341–352. 10.1016/S0969-2126(01)00166-6.

(58) Cusack, S.; Berthet-Colominas, C.; Härtlein, M.; Nassar, N.; Leberman, R. A Second Class of Synthetase Structure Revealed by X-Ray Analysis of Escherichia Coli Seryl-TRNA Synthetase at 2.5 Å. Nature 1990, 347 (6290), 249–255. 10.1038/347249a0.

(59) Grosjean, H.; Nicoghosian, K.; Haumont, E.; Söll, D.; Cedergren, R. Nucleotide Sequences of Two Serine TRNAs with a GGA Anticodon: The Structure-Function Relationships in the Serine Family of E. Coli TRNAs. Nucleic Acids Res. 1985, 13 (15), 5697–5706. 10.1093/nar/13.15.5697.

(60) Gatza, M. L.; Silva, G. O.; Parker, J. S.; Fan, C.; Perou, C. M. An Integrated Genomics Approach Identifies Drivers of Proliferation in Luminal-Subtype Human Breast Cancer. Nat. Genet. 2014, 46 (10), 1051–1059. 10.1038/ng.3073.

(61) Yeon, S. Y.; Jo, Y. S.; Choi, E. J.; Kim, M. S.; Yoo, N. J.; Lee, S. H. Frameshift Mutations in Repeat Sequences of ANK3, HACD4, TCP10L, TP53BP1, MFN1, LCMT2, RNMT, TRMT6, METTL8 and METTL16 Genes in Colon Cancers. Pathol. Oncol. Res. 2018, 24 (3), 617–622. 10.1007/s12253-017-0287-2.

(62) Zhu, X.; Shen, C.; Zhang, W.; Ji, Y.; Xu, S.; Zheng, B.; Chen, Z. Identification and Validation of Prognostic Models and Tumor Microenvironment Infiltration Characteristics for TRNA Modification Regulators in Clear Cell Renal Cell Carcinoma. Oncol. Lett. 2025, 30 (1), 1–17. 10.3892/ol.2025.15108.

(63) Tan, X.-L.; Moyer, A. M.; Fridley, B. L.; Schaid, D. J.; Niu, N.; Batzler, A. J.; Jenkins, G. D.; Abo, R. P.; Li, L.; Cunningham, J. M.; Sun, Z.; Yang, P.; Wang, L. Genetic Variation Predicting Cisplatin Cytotoxicity Associated with Overall Survival in Lung Cancer Patients Receiving Platinum-Based Chemotherapy. Clin. Cancer Res. 2011, 17 (17), 5801–5811. 10.1158/1078-0432.CCR-11-1133.

(64) Hein, M. Y.; Hubner, N. C.; Poser, I.; Cox, J.; Nagaraj, N.; Toyoda, Y.; Gak, I. A.; Weisswange, I.; Mansfeld, J.; Buchholz, F.; Hyman, A. A.; Mann, M. A Human Interactome in Three Quantitative Dimensions Organized by Stoichiometries and Abundances. Cell 2015, 163 (3), 712–723. 10.1016/j.cell.2015.09.053.

(65) Huang, J.; Rauscher, S.; Nawrocki, G.; Ran, T.; Feig, M.; De Groot, B. L.; Grubmüller, H.; MacKerell, A. D. CHARMM36m: An Improved Force Field for Folded and Intrinsically Disordered Proteins. Nat. Methods 2016, 14 (1), 71–73. 10.1038/nmeth.4067.

(66) Abraham, M. J.; Murtola, T.; Schulz, R.; Páll, S.; Smith, J. C.; Hess, B.; Lindahl, E. GROMACS: High Performance Molecular Simulations through Multi-Level Parallelism from Laptops to Supercomputers. SoftwareX 2015, 2, 1–7. 10.1016/j.softx.2015.06.001.

(67) Melo, M. C. R.; Bernardi, R. C.; De La Fuente-Nunez, C.; Luthey-Schulten, Z. Generalized Correlation-Based Dynamical Network Analysis: A New High-Performance Approach for Identifying Allosteric Communications in Molecular Dynamics Trajectories. J. Chem. Phys. 2020, 153 (13), 134104. 10.1063/5.0018980.

(68) Borel, F.; Vincent, C.; Leberman, R.; Härtlein, M. Seryl-TRNA Synthetase from Escherichia Coli: Implication of Its N-Terminal Domain in Aminoacylation Activity and Specificity. Nucleic Acids Res. 1994, 22 (15), 2963–2969. 10.1093/nar/22.15.2963.

(69) Sprinzl, M.; Dank, N.; Nock, S.; Schon, A. Compilation of TRNA Sequences and Sequences of TRNA Genes. Nucleic Acids Res. 1991, 19 (suppl), 2127–2171. 10.1093/nar/19.suppl.2127.

(70) Auffinger, P.; Westhof, E. Singly and Bifurcated Hydrogen-Bonded Base-Pairs in TRNA Anticodon Hairpins and Ribozymes. J. Mol. Biol. 1999, 292 (3), 467–483. 10.1006/JMBI.1999.3080.

(71) Bussi, G.; Laio, A. Using Metadynamics to Explore Complex Free-Energy Landscapes. Nat. Rev. Phys. 2020, 2 (4), 200–212. 10.1038/s42254-020-0153-0.

(72) Tribello, G. A.; Bonomi, M.; Branduardi, D.; Camilloni, C.; Bussi, G. PLUMED 2: New Feathers for an Old Bird. Comput. Phys. Commun. 2014, 185 (2), 604–613. 10.1016/j.cpc.2013.09.018.

(73) Brunk, E.; Rothlisberger, U. Mixed Quantum Mechanical/Molecular Mechanical Molecular Dynamics Simulations of Biological Systems in Ground and Electronically Excited States. Chem. Rev. 2015, 115 (12), 6217–6263. 10.1021/cr500628b.

(74) Branduardi, D.; Gervasio, F. L.; Parrinello, M. From A to B in Free Energy Space. J. Chem. Phys. 2007, 126 (5), 054103. 10.1063/1.2432340.

(75) Craig, D. B.; Arriaga, E. A.; Wong, J. C. Y.; Lu, H.; Dovichi, N. J. Studies on Single Alkaline Phosphatase Molecules: Reaction Rate and Activation Energy of a Reaction Catalyzed by a Single Molecule and the Effect of Thermal Denaturation - The Death of an Enzyme. J. Am. Chem. Soc. 1996, 118 (22), 5245–5253. 10.1021/ja9540839.

(76) Huang, M. H.; Wang, J. T.; Zhang, J. H.; Mao, X. L.; Peng, G. X.; Lin, X.; Lv, D.; Yuan, C.; Lin, H.; Wang, E. D.; Zhou, X. L. Mitochondrial RNA M3C Methyltransferase METTL8 Relies on an Isoform-Specific N-Terminal Extension and Modifies Multiple Heterogenous TRNAs. Sci. Bull. 2023, 68 (18), 2094–2105. 10.1016/j.scib.2023.08.002.

(77) Lentini, J. M.; Alsaif, H. S.; Faqeih, E.; Alkuraya, F. S.; Fu, D. DALRD3 Encodes a Protein Mutated in Epileptic Encephalopathy That Targets Arginine TRNAs for 3-Methylcytosine Modification. Nat. Commun. 2020 111 2020, 11 (1), 1–14. 10.1038/s41467-020-16321-6.

(78) Lentini, J. M.; Bargabos, R.; Chen, C.; Fu, D. Methyltransferase METTL8 Is Required for 3-Methylcytosine Modification in Human Mitochondrial TRNAs. J. Biol. Chem. 2022, 298 (4), 101788. 10.1016/j.jbc.2022.101788.

(79) Liu, J.; Yue, Y.; Han, D.; Wang, X.; Fu, Y.; Zhang, L.; Jia, G.; Yu, M.; Lu, Z.; Deng, X.; Dai, Q.; Chen, W.; He, C. A METTL3-METTL14 Complex Mediates Mammalian Nuclear RNA N6-Adenosine Methylation. Nat. Chem. Biol. 2014, 10 (2), 93–95. 10.1038/nchembio.1432.

(80) Wang, P.; Doxtader, K. A.; Nam, Y. Structural Basis for Cooperative Function of Mettl3 and Mettl14 Methyltransferases. Mol. Cell 2016, 63 (2), 306–317. 10.1016/j.molcel.2016.05.041.

(81) Œledź, P.; Jinek, M. Structural Insights into the Molecular Mechanism of the m(6)A Writer Complex. Elife 2016, 5, e18434. 10.7554/eLife.18434.

(82) Li, J.; Wang, L.; Hahn, Q.; Nowak, R. P.; Viennet, T.; Orellana, E. A.; Roy Burman, S. S.; Yue, H.; Hunkeler, M.; Fontana, P.; Wu, H.; Arthanari, H.; Fischer, E. S.; Gregory, R. I. Structural Basis of Regulated M7G TRNA Modification by METTL1–WDR4. Nature 2023, 613 (7943), 391–397. 10.1038/s41586-022-05566-4.

(83) Ruiz-Arroyo, V. M.; Raj, R.; Babu, K.; Onolbaatar, O.; Roberts, P. H.; Nam, Y. Structures and Mechanisms of TRNA Methylation by METTL1–WDR4. Nature 2023, 613 (7943), 383–390. 10.1038/s41586-022-05565-5.

(84) Ishiguro, K.; Fujimura, A.; Shirouzu, M. Structural Insights into TRNA Recognition of the Human FTSJ1-THADA Complex. *Commun*. Biol. 2025, 8 (1), 1–11. 10.1038/s42003-025-08278-3.

(85) Xu, H.; Shi, R.; Han, W.; Cheng, J.; Xu, X.; Cheng, K.; Wang, L.; Tian, B.; Zheng, L.; Shen, B.; Hua, Y.; Zhao, Y. Structural Basis of 5′ Flap Recognition and Protein–Protein Interactions of Human Flap Endonuclease 1. Nucleic Acids Res. 2018, 46 (21), 11315–11325. 10.1093/NAR/GKY911.

(86) Ali, R. H.; Orellana, E. A.; Lee, S. H.; Chae, Y.-C.; Chen, Y.; Clauwaert, J.; Kennedy, A. L.; Gutierrez, A. E.; Papke, D. J.; Valenzuela, M.; Silverman, B.; Falzetta, A.; Ficarro, S. B.; Marto, J. A.; Fletcher, C. D. M.; Perez-Atayde, A.; Alcindor, T.; Shimamura, A.; Prensner, J. R.; Gregory, R. I.; Gutierrez, A. A Methyltransferase-Independent Role for METTL1 in TRNA Aminoacylation and Oncogenic Transformation. Mol. Cell 2025, 85 (5), 948–961.e11. 10.1016/j.molcel.2025.01.003.

(87) Almatarneh, M. H.; Kayed, G. G.; Altarawneh, M.; Zhao, Y.; Verma, A. Computational Insights in DNA Methylation: Catalytic and Mechanistic Elucidations for Forming 3-Methyl Cytosine. J. Chem. 2022, 2022 (1), 2673396. 10.1155/2022/2673396.

(88) Dukatz, M.; Requena, C. E.; Emperle, M.; Hajkova, P.; Sarkies, P.; Jeltsch, A. Mechanistic Insights into Cytosine-N3 Methylation by DNA Methyltransferase DNMT3A. J. Mol. Biol. 2019, 431 (17), 3139–3145. 10.1016/j.jmb.2019.06.015.

(89) Corbeski, I.; Vargas-Rosales, P. A.; Bedi, R. K.; Deng, J.; Coelho, D.; Braud, E.; Iannazzo, L.; Li, Y.; Huang, D.; Ethève-Quelquejeu, M.; Cui, Q.; Caflisch, A. The Catalytic Mechanism of the RNA Methyltransferase METTL3. Elife 2024, 12. 10.7554/ELIFE.92537.

(90) Liu, R. J.; Long, T.; Li, J.; Li, H.; Wang, E. D. Structural Basis for Substrate Binding and Catalytic Mechanism of a Human RNA:M5C Methyltransferase NSun6. Nucleic Acids Res. 2017, 45 (11), 6684–6697. 10.1093/NAR/GKX473.

(91) Lee, B. W. L.; Chuah, Y. H.; Yoon, J.; Grinchuk, O. V.; Liang, Y.; Hirpara, J. L.; Shen, Y.; Wang, L. C.; Lim, Y. T.; Zhao, T.; Sobota, R. M.; Yeo, T. T.; Wong, A. L. A.; Teo, K.; Nga, V. D. W.; Tan, B. W. Q.; Suda, T.; Toh, T. B.; Pervaiz, S.; Lin, Z.; Ong, D. S. T. METTL8 Links Mt-TRNA M3C Modification to the HIF1α/RTK/Akt Axis to Sustain GBM Stemness and Tumorigenicity. Cell Death Dis. 2024, 15 (5), 338. 10.1038/s41419-024-06718-2.

(92) Wu, E. L.; Cheng, X.; Jo, S.; Rui, H.; Song, K. C.; Dávila-Contreras, E. M.; Qi, Y.; Lee, J.; Monje-Galvan, V.; Venable, R. M.; Klauda, J. B.; Im, W. CHARMM-GUI Membrane Builder toward Realistic Biological Membrane Simulations. J. Comput. Chem. 2014, 35 (27), 1997–2004. 10.1002/jcc.23702.

(93) Xu, Y.; Vanommeslaeghe, K.; Aleksandrov, A.; MacKerell, A. D.; Nilsson, L. Additive CHARMM Force Field for Naturally Occurring Modified Ribonucleotides. J. Comput. Chem. 2016, 37 (10), 896–912. 10.1002/JCC.24307.

(94) Jorgensen, W. L.; Chandrasekhar, J.; Madura, J. D.; Impey, R. W.; Klein, M. L. Comparison of Simple Potential Functions for Simulating Liquid Water. J. Chem. Phys. 1983, 79 (2), 926–935. 10.1063/1.445869.

(95) Love, O.; Galindo-Murillo, R.; Roe, D. R.; Dans, P. D.; Cheatham, T. E.; Bergonzo, C. ModXNA: A Modular Approach to Parametrization of Modified Nucleic Acids for Use with Amber Force Fields. J. Chem. Theory Comput. 2024, 20 (21), 9354–9363. 10.1021/acs.jctc.4c01164.

(96) Wang, J.; Wang, W.; Kollman, P. A.; Case, D. A. Antechamber, An Accessory Software Package For Molecular Mechanical Calculations. J. Am. Chem. Soc 2001, 222, U403.

(97) Li, P.; Song, L. F.; Merz, K. M. Systematic Parameterization of Monovalent Ions Employing the Nonbonded Model. J. Chem. Theory Comput. 2015, 11 (4), 1645–1657. 10.1021/ct500918t.

(98) Essmann, U.; Perera, L.; Berkowitz, M. L.; Darden, T.; Lee, H.; Pedersen, L. G. A Smooth Particle Mesh Ewald Method. J. Chem. Phys. 1995, 103 (19), 8577–8593. 10.1063/1.470117.

(99) Bussi, G.; Donadio, D.; Parrinello, M. Canonical Sampling through Velocity Rescaling. J. Chem. Phys. 2007, 126 (1), 014101. 10.1063/1.2408420.

(100) Parrinello, M.; Rahman, A. Polymorphic Transitions in Single Crystals: A New Molecular Dynamics Method. J. Appl. Phys. 1981, 52 (12), 7182–7190. 10.1063/1.328693.

(101) Michaud-Agrawal, N.; Denning, E. J.; Woolf, T. B.; Beckstein, O. MDAnalysis: A Toolkit for the Analysis of Molecular Dynamics Simulations. J. Comput. Chem. 2011, 32 (10), 2319–2327. 10.1002/jcc.21787.

(102) Bouysset, C.; Fiorucci, S. ProLIF: A Library to Encode Molecular Interactions as Fingerprints. J. Cheminform. 2021, 13 (1), 72. 10.1186/s13321-021-00548-6.

(103) Barducci, A.; Bussi, G.; Parrinello, M. Well-Tempered Metadynamics: A Smoothly Converging and Tunable Free-Energy Method. Phys. Rev. Lett. 2008, 100 (2), 020603. 10.1103/PhysRevLett.100.020603.

(104) Hutter, J.; Iannuzzi, M.; Schiffmann, F.; Vandevondele, J. Cp2k: Atomistic Simulations of Condensed Matter Systems. Wiley Interdiscip. Rev. Comput. Mol. Sci. 2014, 4 (1), 15–25. 10.1002/wcms.1159.

(105) Becke, A. D. Density-Functional Exchange-Energy Approximation with Correct Asymptotic Behavior. Phys. Rev. A 1988, 38 (6), 3098–3100. 10.1103/PhysRevA.38.3098.

(106) Lee, C.; Yang, W.; Parr, R. G. Development of the Colle-Salvetti Correlation-Energy Formula into a Functional of the Electron Density. Phys. Rev. B 1988, 37 (2), 785–789. 10.1103/PhysRevB.37.785.

(107) VandeVondele, J.; Hutter, J. Gaussian Basis Sets for Accurate Calculations on Molecular Systems in Gas and Condensed Phases. J. Chem. Phys. 2007, 127 (11), 1–10. 10.1063/1.2770708.

(108) Grimme, S.; Antony, J.; Ehrlich, S.; Krieg, H. A Consistent and Accurate Ab Initio Parametrization of Density Functional Dispersion Correction (DFT-D) for the 94 Elements H-Pu. J. Chem. Phys. 2010, 132 (15), 154104. 10.1063/1.3382344.

(109) Torrie, G. M.; Valleau, J. P. Nonphysical Sampling Distributions in Monte Carlo Free-Energy Estimation: Umbrella Sampling. J. Comput. Phys. 1977, 23 (2), 187–199. 10.1016/0021-9991(77)90121-8.

(110) Hub, J. S.; De Groot, B. L.; Van Der Spoel, D. G-Whams-a Free Weighted Histogram Analysis Implementation Including Robust Error and Autocorrelation Estimates. J. Chem. Theory Comput. 2010, 6 (12), 3713–3720. 10.1021/ct100494z.

(111) Wang, J. X.; Lee, E. R.; Morales, D. R.; Lim, J.; Breaker, R. R. Riboswitches That Sense S-Adenosylhomocysteine and Activate Genes Involved in Coenzyme Recycling. Mol. Cell 2008, 29 (6), 691–702. 10.1016/j.molcel.2008.01.012.

(112) Pham, H.; Kumar, M.; Martinez, A. R.; Ali, M.; Lowery, R. G. Development and Validation of a Generic Methyltransferase Enzymatic Assay Based on an SAH Riboswitch. SLAS Discov. Adv. life Sci. R D 2024, 29 (4). 10.1016/J.SLASD.2024.100161.

(113) Nidoieva, Z.; Sabin, M. O.; Dewald, T.; Weldert, A. C.; Hoba, S. N.; Helm, M.; Barthels, F. A Microscale Thermophoresis-Based Enzymatic RNA Methyltransferase Assay Enables the Discovery of DNMT2 Inhibitors. Commun. Chem. 2025 81 2025, 8 (1), 1–9. 10.1038/s42004-025-01439-9.

