## Supporting Figs. S1-S17, Supporting Table S1 for "Recognition and catalytic mechanism of tRNA m^3^C methyltransferase METTL6"

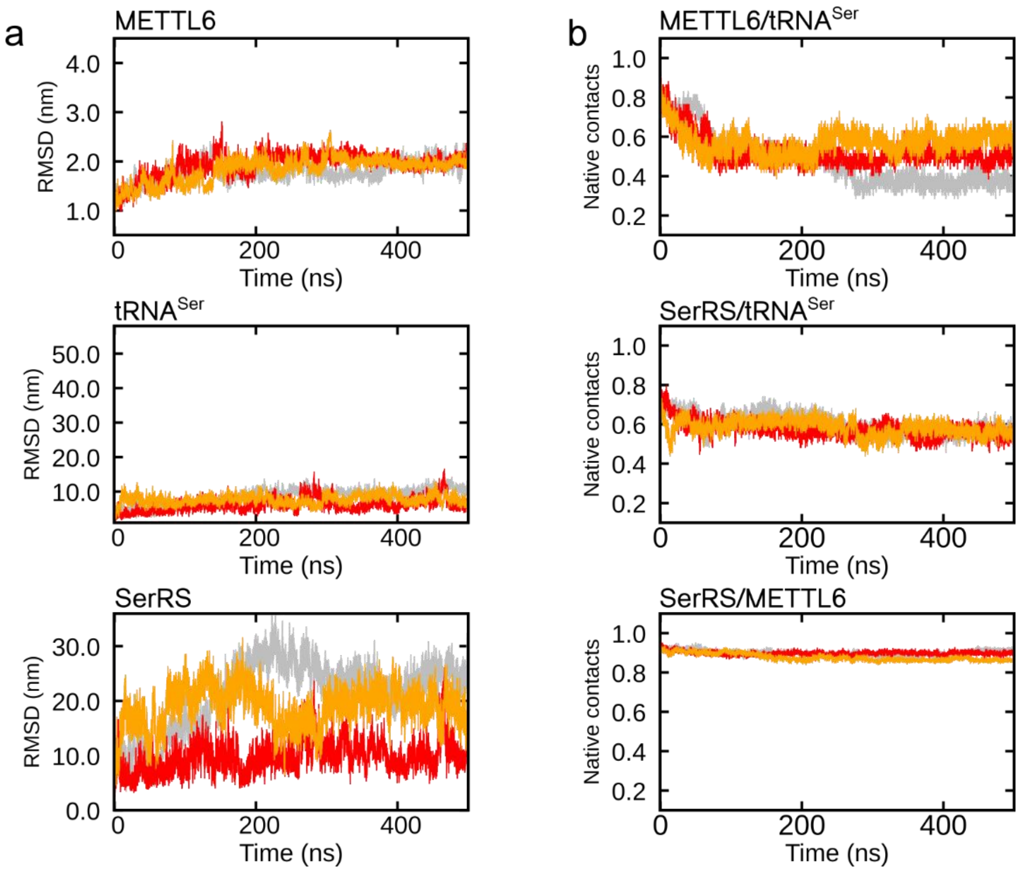


**Figure S1**. **METTL6–SerRS–tRNA$^{Ser}$ complex remained well-folded during MD simulations.** MD simulations were performed in triplicates, with each replica represented by a different line color. While simulations were performed with the complex exhibiting 1:2:2 stoichiometry (i.e. 1 METTL6, 2 SerRS, 2 tRNA$^{Ser}$ molecules) stoichiometry, results are shown only for SerRS and tRNA$^{Ser}$ molecules in contact with METTL6. (a) Backbone relative mean square deviation (RMSD) as a function of simulation time. Frames were superimposed on the METTL6 protein backbone. (b) Fraction of intermolecular native contacts present at different interfaces in the METTL6–SerRS–tRNA$^{Ser}$ complex as a function of simulation time. The cryo-EM structure (PDBID: 8P7B) was used for calculation of native contacts. Two residues were considered in contact if the distance between their Cα atoms (or Cα and P atoms in case of interfaces involving tRNA$^{Ser}$) was below 9 Å.

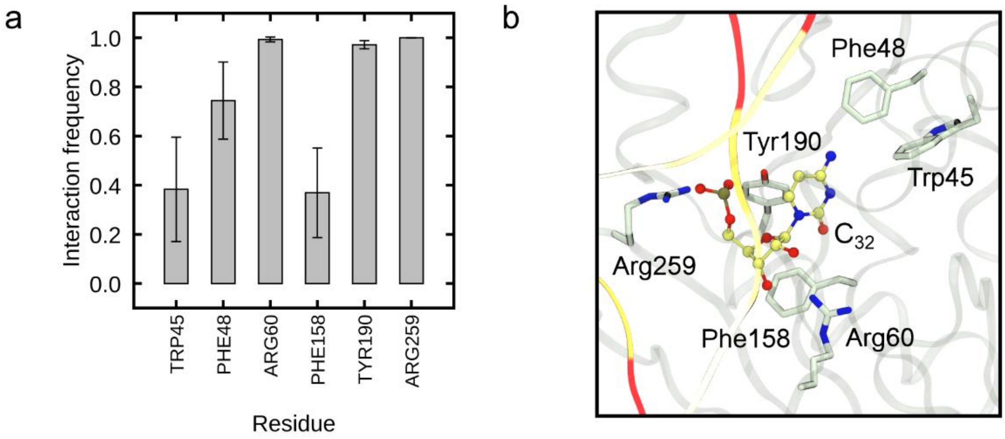


**Figure S2. METTL6 residues stabilizing the reactive conformation of $C_{32}$ in the catalytic cavity as observed during MD simulations of the METTL6–SerRS–tRNA$^{Ser}$ complex.** (a) Interaction frequency for METTL6 residues interacting with $C_{32}$ for at least 30 % of the simulation time. Error bars denote ± s.d. from the mean for $n$ = 3 simulation replicas. (b) Snapshot of the catalytic cavity. METTL6 residues listed in panel (a) are shown as licorice, while $C_{32}$ is shown in CPK representation.

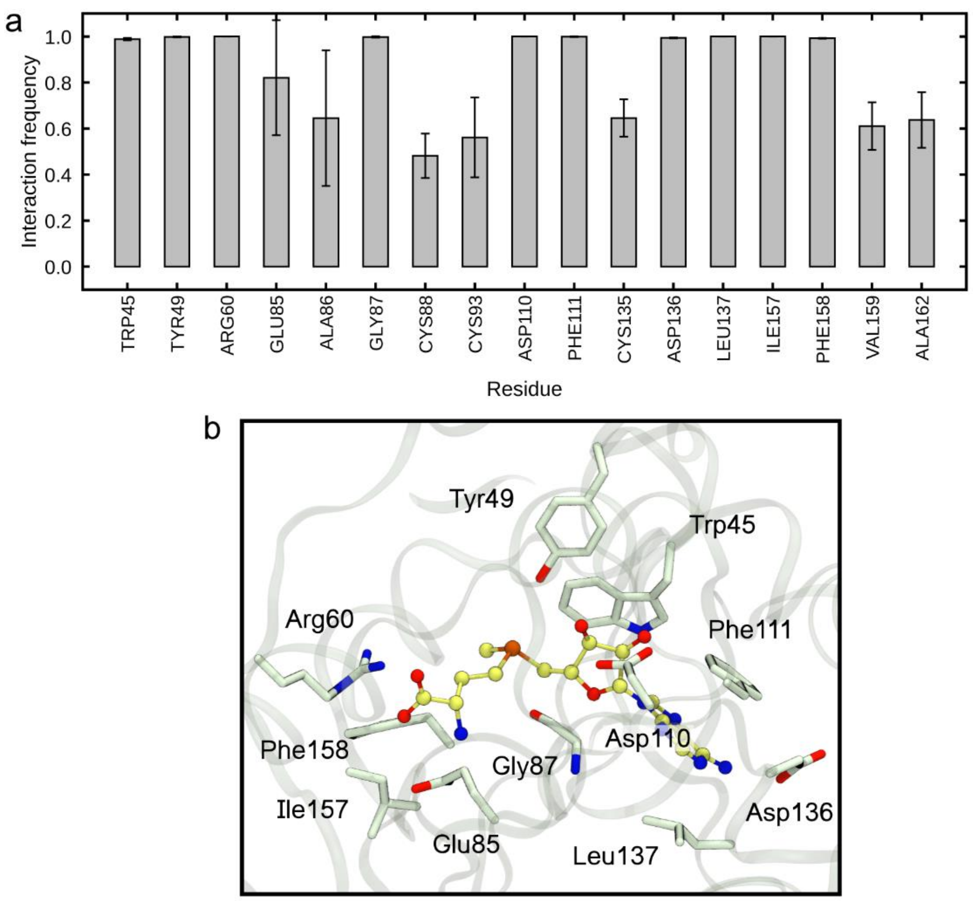


**Figure S3. METTL6 residues stabilizing the reactive conformation of SAM in the catalytic cavity as observed during MD simulations of the METTL6–SerRS–tRNA$^{Ser}$ complex.** (a) Interaction frequencies between METTL6 residues and $C_{32}$. The depicted residues interact for at least 30 % of the simulation time. Error bars denote ± s.d. from the mean for $n$ = 3 simulation replicas. (b) Snapshot of the catalytic cavity. METTL6 residues listed in panel (a) are shown as licorice, while SAM is shown in CPK representation.

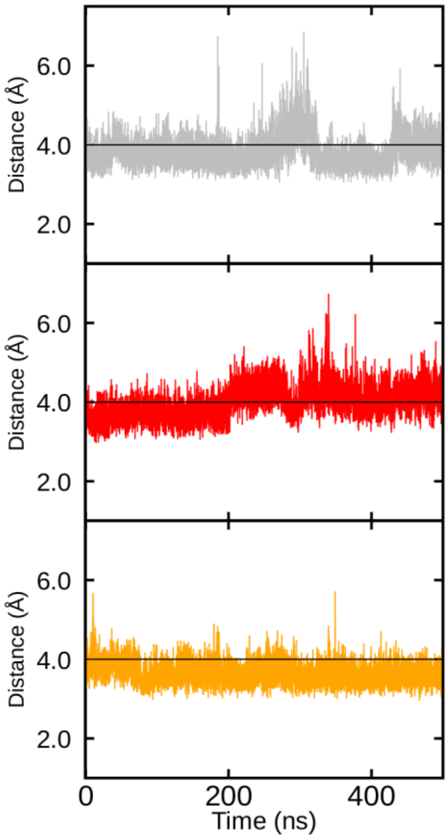


**Figure S4. Distance between the $C_{32}$:N3-SAM:$CH_3$ atoms as observed during MD simulations of the METTL6–SerRS–tRNA$^{Ser}$ complex.** Each panel corresponds to a single replica. The horizontal line at 4 Å represent the threshold for differentiating reactive and non-reactive conformations.

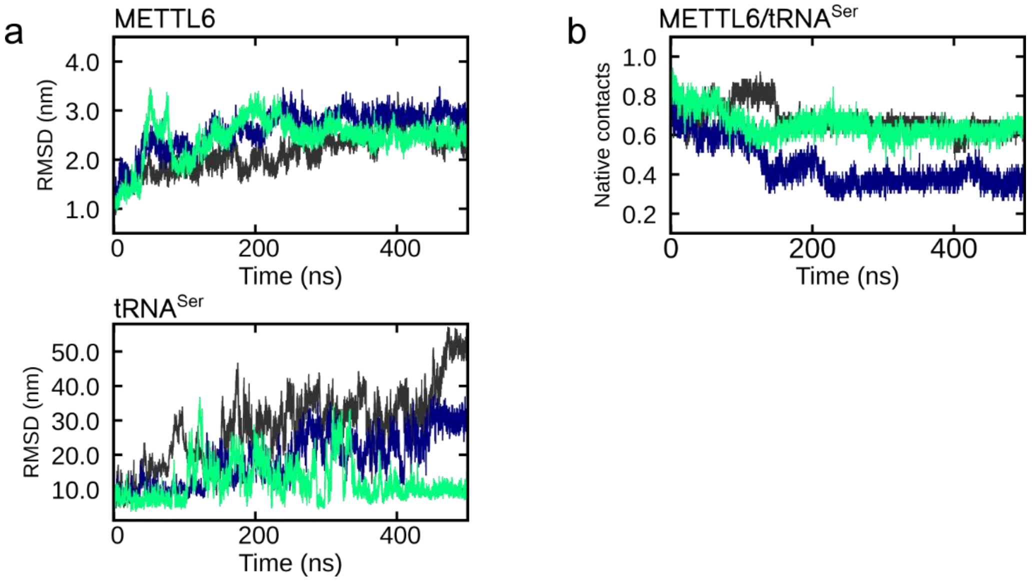


**Figure S5. Stability of the METTL6–tRNA$^{Ser}$ complex during MD simulations.** MD simulations were performed in triplicates, with each replica represented by a different line color. (a) Backbone relative mean square deviation (RMSD) as a function of simulation time. Frames were superimposed on the METTL6 protein backbone. (b) Fraction of intermolecular native contacts present at the METTL6/tRNA$^{Ser}$ interface as a function of simulation time. The cryo-EM structure (PDBID: 8P7B) was used for calculation of native contacts. Two residues were considered in contact if the distance between their Cα and P atoms was below 9 Å.

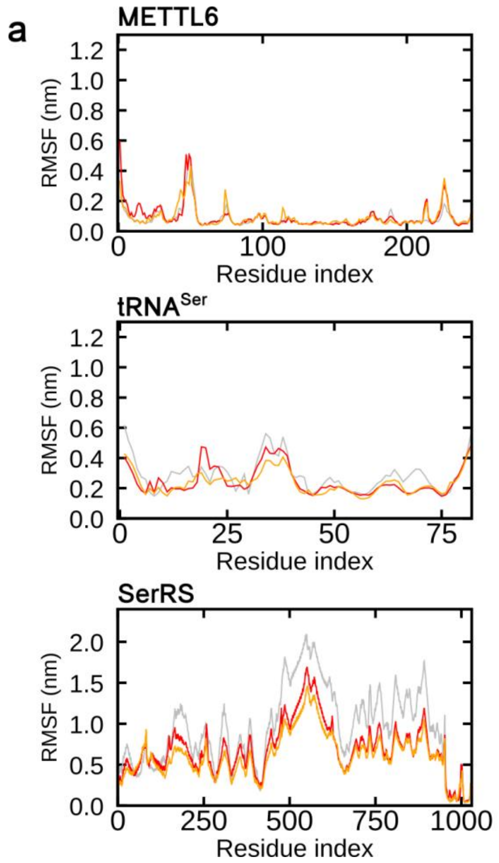


**Figure S6. Relative mean square fluctuation (RMSF) per residue for each molecular unit.** (a) RMSF as observed during MD simulations of the 1:2:2 METTL6-SerRS-tRNA$^{Ser}$ complex. RMSF values were calculated from the positions of Cα or P atoms along the trajectory. MD simulations were performed in triplicates, with each replica represented by a different line color. Results are shown only for SerRS and tRNA$^{Ser}$ molecules in contact with METTL6. (b) RMSF as observed during MD simulations of the METTL6-tRNA$^{Ser}$ complex in the absence of SerRS.

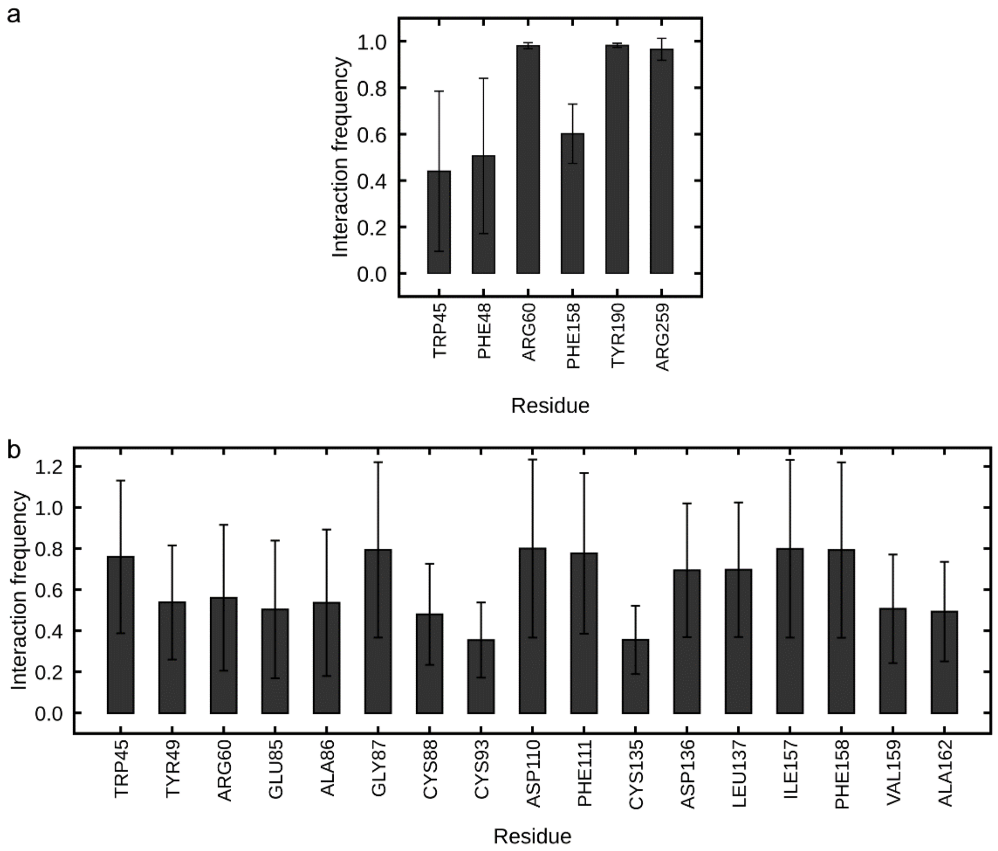


**Figure S7. Interaction of METTL6 residues in the catalytic cavity with $C_{32}$ and SAM as observed during MD simulations of the METTL6–tRNA$^{Ser}$ complex in the absence of SerRS.** (a) Interaction frequency for METTL6 residues interacting with $C_{32}$ for at least 30 % of the simulation time. (b) Interaction frequency for METTL6 residues interacting with SAM for at least 30 % of the simulation time. Error bars denote ± s.d. from the mean for $n$ = 3 simulation replicas.

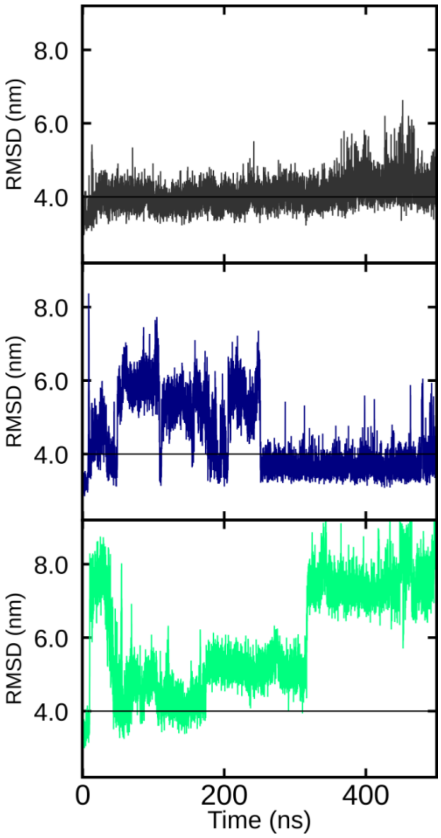


**Figure S8. Distance between the $C_{32}$:N3-SAM:$CH_3$ atoms as observed during MD simulations of the METTL6–$tRNA^{Ser}$ complex in the absence of SerRS.** Each panel corresponds to a single replica. The horizontal line at 4 Å represent the threshold for differentiating reactive and non-reactive conformations.

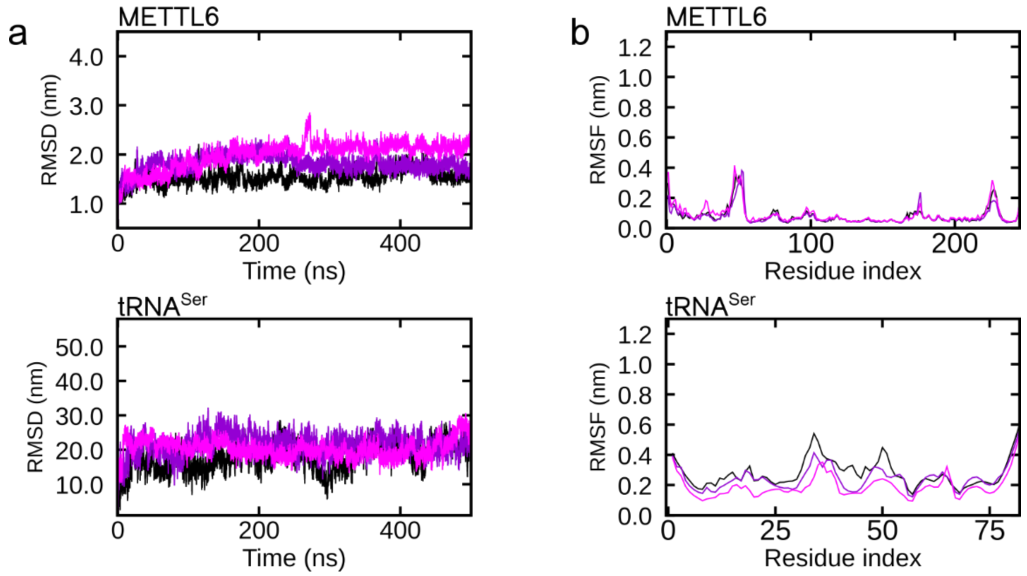


**Figure S9. Stability of the METTL6–tRNA$^{Ser}$ complex during MD simulations performed with the Amber forcefields.** MD simulations were performed in triplicates, with each replica represented by a different line color. (a) Backbone relative mean square deviation (RMSD) as a function of simulation time. Frames were superimposed on the METTL6 protein backbone. (b) Relative mean square fluctuation (RMSF) per residue. RMSF values were calculated from the positions of Cα or P atoms along the trajectory.

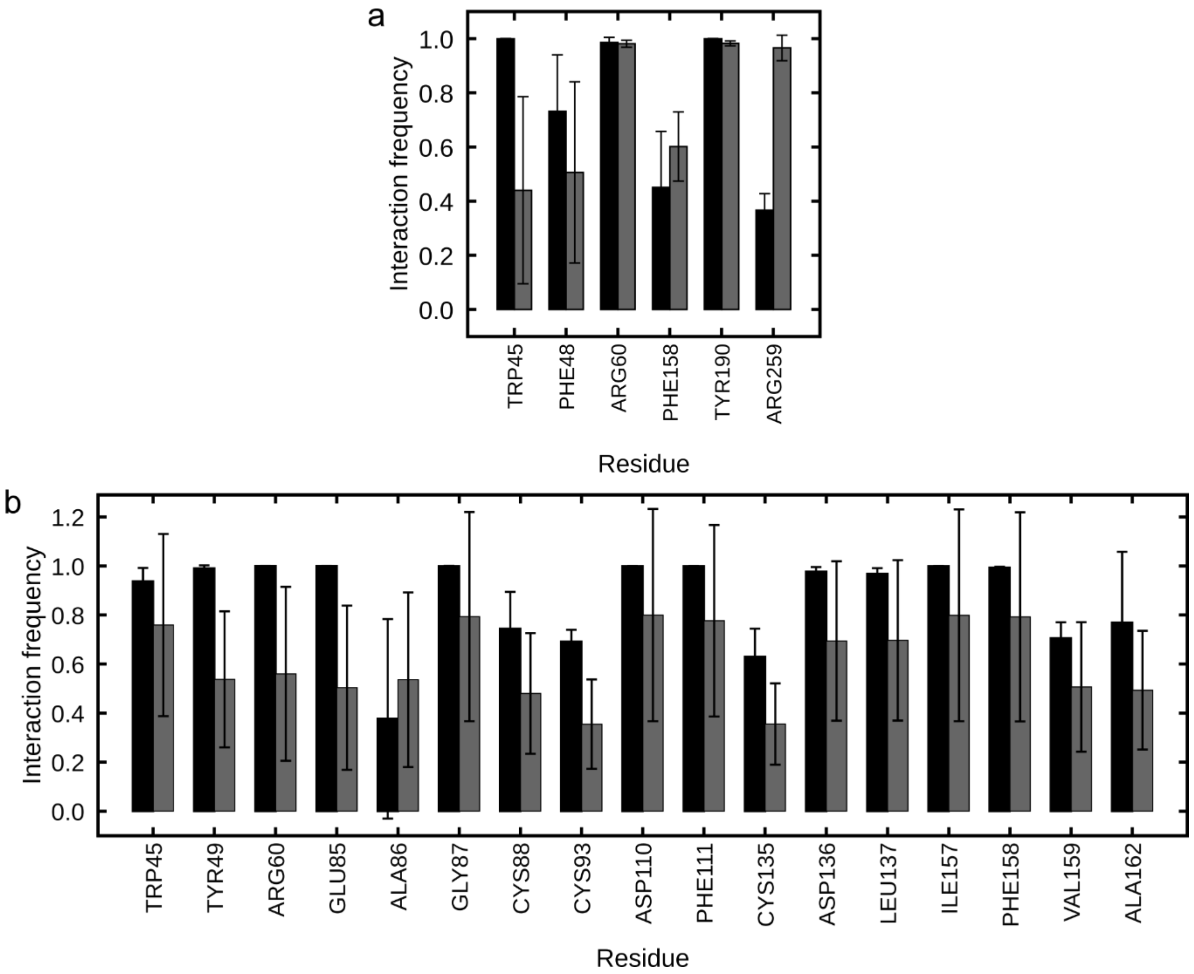


**Figure S10. Influence of force field selection on the frequency of interactions of METTL6 residues with $C_{32}$ and SAM during MD simulations of the METTL6–tRNA$^{Ser}$ complex in the absence of SerRS.** Black bars denote the results obtained with the Amber force field, while dark gray bars correspond to simulations performed with the CHARMM force field. (a) Interaction frequency for METTL6 residues interacting with $C_{32}$. (b) Interaction frequency for METTL6 residues interacting with SAM. Error bars denote ± s.d. from the mean for $n$ = 3 simulation replicas.

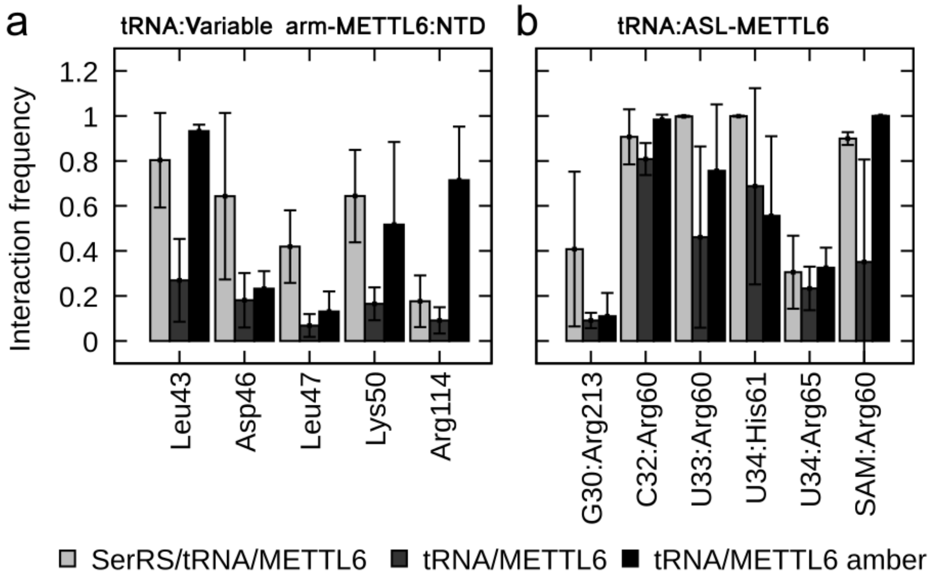


**Figure S11. Influence of force field selection on the frequency of interactions between METTL6 and tRNA$^{Ser}$.** Light gray bars show results obtained for the METTL6-SerRS-tRNA$^{Ser}$ complex with the CHARMM force field, dark gray bars correspond to simulations performed for the METTL6-tRNA$^{Ser}$ complex with the CHARMM force field, while black bars denote the results obtained for the METTL6-tRNA$^{Ser}$ complex with the Amber force field. (a) Interaction frequency between the METTL6 N-terminal domain (NTD) and tRNA$^{Ser}$ variable arm. (b) Interaction frequency for hydrogen bonding between METTL6 and tRNA$^{Ser}$ anticodon stem loop (ASL). Error bars denote ± s.d. from the mean for $n$ = 3 simulation replicas.

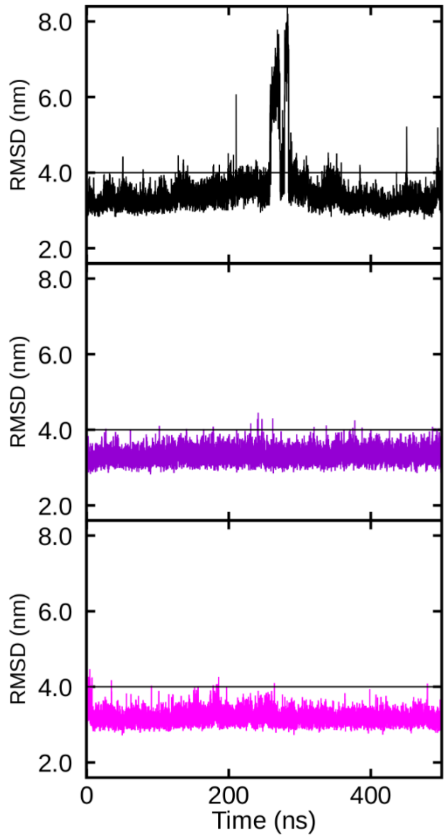


**Figure S12. Distance between the $C_{32}$:N3-SAM:$CH_3$ atoms as observed during control MD simulations performed with the Amber force field.** Simulations were performed for the METTL6–tRNA$^{Ser}$ complex in the absence of SerRS. Each panel corresponds to a single replica. The horizontal line at 4 Å represent the threshold for differentiating reactive and non-reactive conformations.

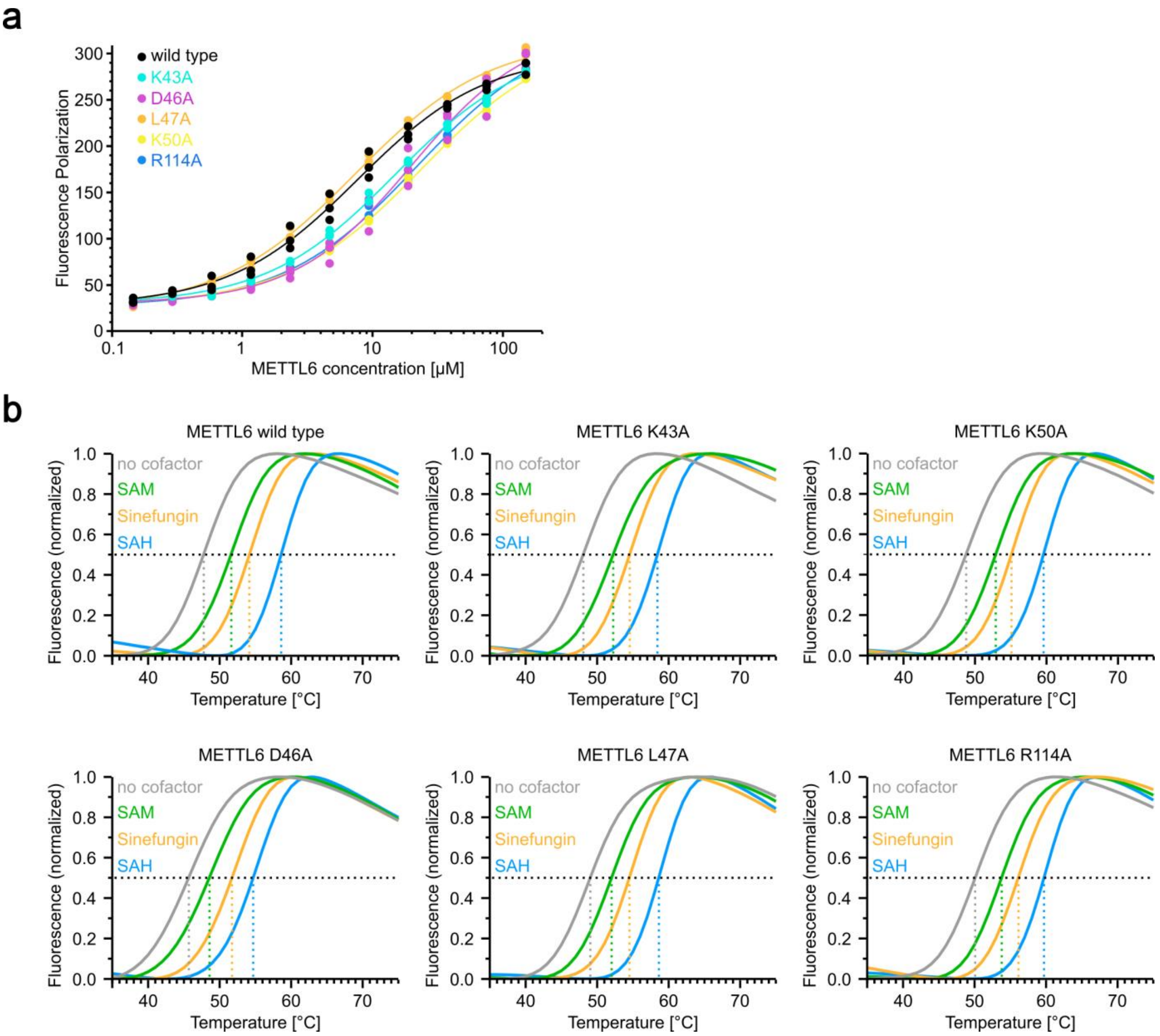


**Figure S13. Biochemical characterization of METTL6 interaction with tRNA$^{Ser}$ and SAM analogues.** (a) *In vitro* interaction of different METTL6 variants with tRNA$^{Ser}$ determined by fluorescence polarization. Normalized sigmoidal fits of three replicates are shown with each replicate indicated as an individual datapoint. (b) *In vitro* thermal stability assay of METTL6 variants in the absence and presence of different cofactors SAM, SAH and sinefungin. A representative melting curve out of at least two replicates is shown for each METTL6 variant.

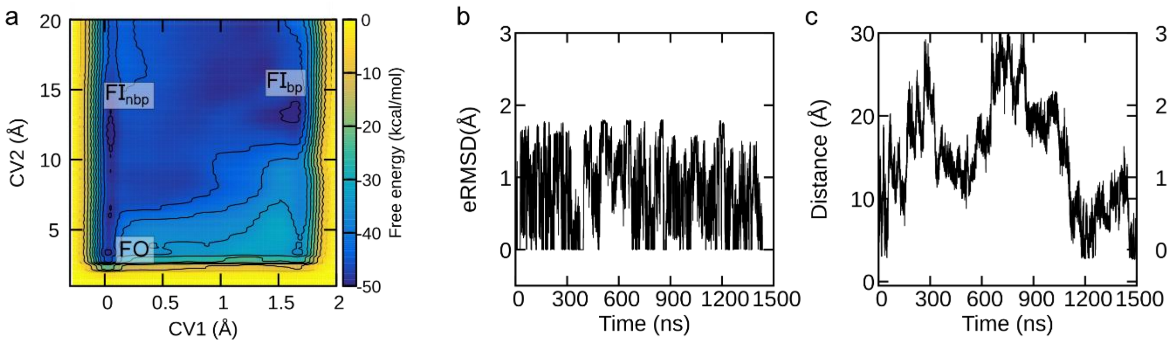


**Figure S14. Metadynamics simulations of the tRNA$^{Ser}$ anticodon remodeling.** Collective variable 1 (CV1) corresponds to eRMSD, while CV2 reports the distance between $C_{32}$:N3 and SAM:$CH_3$ atoms. Prior to METTL6 binding, $C_{32}$ is flipped-in towards the anticodon stem loop and forms a non-canonical base pair with $A_{38}$ ($FI_{bp}$ state). In the reactive state, reached upon METTL6 binding, $C_{32}$ assumes a flipped-out conformation (FO state). Our metadynamics simulation uncovered an additional on path intermediate conformation, where $C_{32}$ is flipped-in, but does not base pair with $A_{38}$ ($FI_{nbp}$ state). (a) Two-dimensional free energy surface of the tRNA$^{Ser}$ anticodon stem loop conformational transition from the $FI_{bp}$ to the FO state. (b) Value of eRMSD as a function of simulation time. (c) Value of the distance between $C_{32}$:N3 and SAM:$CH_3$ atoms as a function of simulation time.

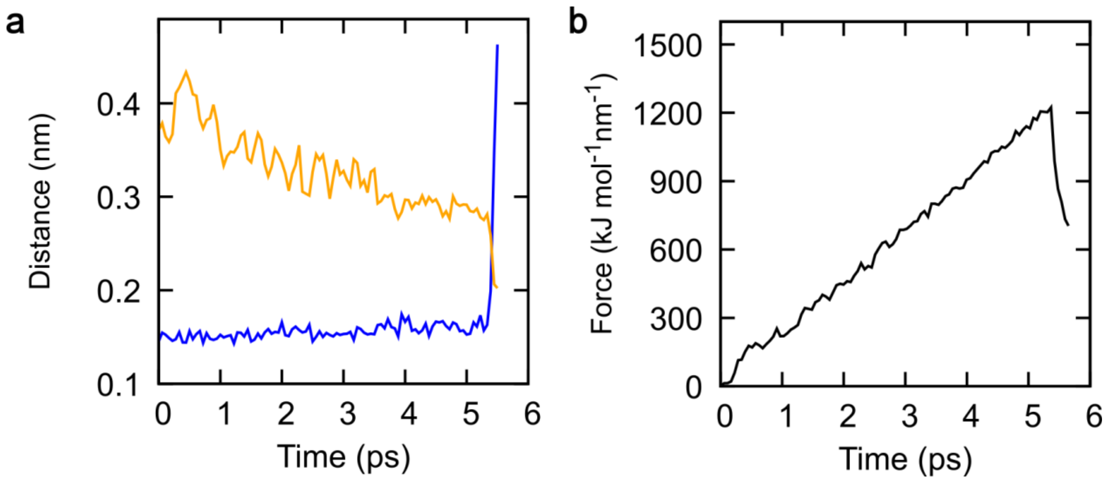


**Figure S15. Steered hybrid quantum-classical (QM/MM) molecular dynamics (MD) simulations of the $C_{32}$ methylation reaction.** The reaction simulation was started from the product state with $m^3C_{32}$ already formed, since this state was experimentally resolved. Reaction coordinate was defined as the difference in length of the forming ($m^3C_{32}$:$CH_3$-SAH:S) and breaking ($m^3C_{32}$:N3-$m^3C_{32}$:$CH_3$) bond. The reaction was observed around 5 ps, suggesting the selected reaction coordinate is appropriate for performing more accurate enhanced sampling simulations. (a) Distance between $m^3C_{32}$:N3 and $m^3C_{32}$:$CH_3$ atoms as a function of time (blue line) and the distance between $m^3C_{32}$:$CH_3$ and SAH:S atoms (orange line). (b) Pulling force applied to the reaction coordinate as a function of simulation time.

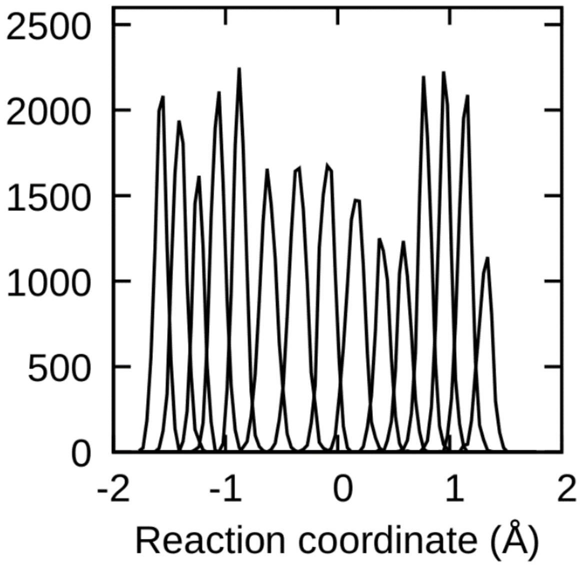


**Figure S16. WHAM histogram depicting the sampling of the reaction coordinate in different simulation windows during QM/MM umbrella sampling simulations.**

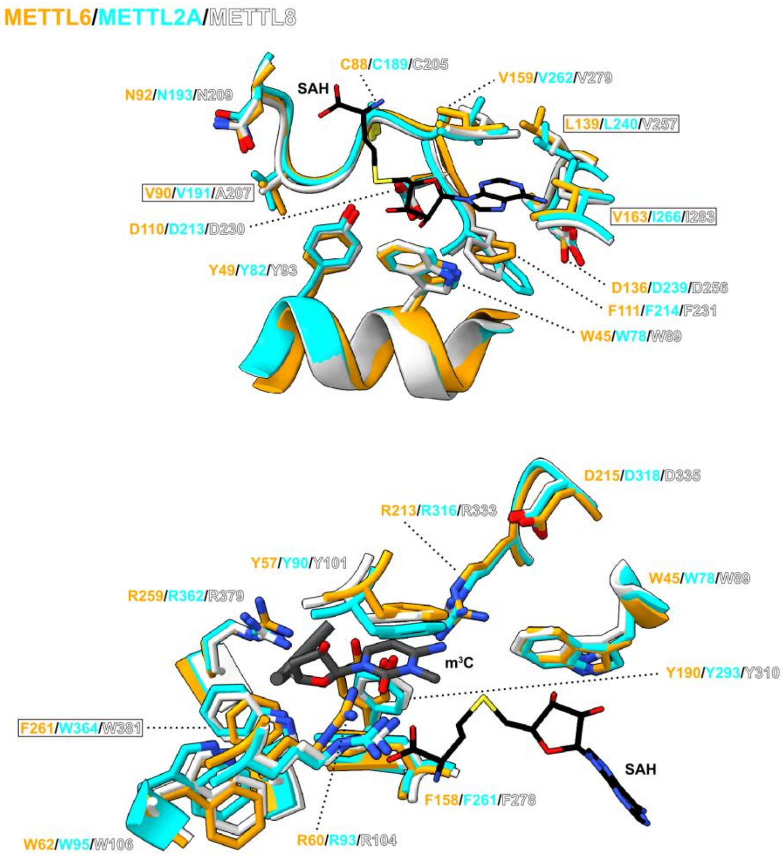


**Figure S17. Active site of human m$^3$C RNA methyltransferases is highly conserved.** Residues interacting with SAH and m$^3$C$_{32}$ are shown as licorice in orange, cyan and white for METTL2, METTL6 and METTL8, respectively. SAH and m$^3$C$_{32}$ are shown in gray.

**Table S1. List of aptamer sequences.**

| Name | Sequence |
|---|---|
| P1 | Biotin-AGGGUCUGCCGAGGAGCGCUGCGACCCUUUAAUUCGGGGGCCAGGC UCGGCAAUGAUGCC |
| P2 | AUGAUCAACGGCGCUCGC-Cy5 |
